# UBE3C retrofits the proteasome to enforce degradation of ultra-stable folds

**DOI:** 10.64898/2026.02.24.707655

**Authors:** Shitao Zou, Deyao Yin, Miyun Shi, Enbo Chen, Xin Luo, Matthew R Baird, Lihong Zhao, Shuo Cao, Di Wu, Shuwen Zhang, Daniel Finley, Youdong Mao

## Abstract

Neurodegeneration and proteinopathies arise from mutated or misfolded proteins that form ultra-stable, proteotoxic aggregates that evade proteasomal degradation despite ubiquitylation^1–8^. These conditions are exacerbated by the deubiquitylase USP14 that can prematurely rescue substrates^9–11^. To counter these obstacles to effective proteolysis, eukaryotes evolved the highly conserved HECT-type E3 ligase UBE3C—dysregulated in neurodegeneration^12,13^ and overexpressed in cancers^14–17^—to reprogram the 26S proteasome toward heightened activity via unknown mechanisms^18–25^. Here we visualized functional dynamics of the human UBE3C-retrofitted proteasome enforcing degradation of a re-engineered superfolder GFP that normally escapes proteolysis, using time-resolved cryo-electron microscopy. A continuum of non-equilibrium conformations, comprising fourteen proteasome conformers orthogonally combined with four UBE3C states or three USP14 states, reveals a cryptic UBE3C-receptor site in the proteasomal lid and captures key intermediates of ubiquitin-chain elongation and branching at four linkage-specific ubiquitin-binding sites. Remarkably, UBE3C creates an extreme shortcut for ubiquitin shuttling that bypasses USP14, promotes USP14 recycling and allosterically strengthens substrate-unfolding forces of the AAA-ATPase unfoldase to ultimately surmount the unfolding energy barrier of ultra-stable proteins. These findings define the complete functional cycle of the UBE3C-reprogrammed proteasome, illuminating how UBE3C simultaneously controls USP14 and the proteasome to enforce clearance of ultra-stable folds, and establish crucial mechanistic foundations for proteolysis-targeting therapeutic discovery.

---

Ultra-stable protein structures—including misfolded or mutated protein aggregates, and hyperthermophile-expressed proteins that resist unfolding and proteolytic degradation even after ubiquitylation—can trigger severe cellular toxicity by disrupting key homeostatic pathways and overwhelming proteostasis networks^1,2^. The ubiquitin-proteasome system (UPS) is the central proteolytic machinery for coping with such proteotoxic stress and regulates nearly every aspect of intracellular activities in eukaryotes^26–28^. UPS dysregulation is a hallmark of neurodegeneration, ageing and cancer^1–8^. In frontotemporal dementia and amyotrophic lateral sclerosis, hyper-stable poly-GA or TDP-43 aggregates trap the 26S proteasome in substrate-engaged conformations^3,4,10^, thereby impairing substrate turnover, exacerbating proteotoxic stress and accelerating disease progression^5,6^. To act upon these ultra-stable misfolds, the 26S proteasome reversibly recruits specialized enzymatic factors, such as the highly conserved HECT-type E3 ligase UBE3C (RAUL/KIAA10), forming higher-molecular-weight supercomplexes with enhanced unfoldase activity and increased degradative processivity that are assembled in a stress-inducible manner to clear ultra-stable, otherwise refractory substrates and maintain protein quality under tight control^20,29–33^.

Among the hundreds of human E3 ubiquitin ligases, UBE3C (and its yeast ortholog Hul5) is the only processivity factor known to enhance proteasomal degradation through association with the proteasome^18–25,34–36^. UBE3C catalyzes the synthesis of K48-, K29- or K33-linked ubiquitin chains, driving complete breakdown of substrates that otherwise stall the proteasome or resist unfolding^12,13,21,35,37^. UBE3C is dysregulated in neurodegenerative disease^12,13^ and is overexpressed in multiple cancers, including hepatocellular carcinoma, gastric cancer, non-small-cell lung cancer and breast cancer^14–17^, underscoring its central role in preserving proteome fidelity and its potential as a therapeutic target. The deubiquitylating (DUB) enzyme USP14, also implicated in neurodegeneration and cancer^38–42^, suppresses the proteasomal degradation by prematurely trimming ubiquitin chains and is antagonized by UBE3C, yet paradoxically promotes UBE3C recruitment to the proteasome and co-cycles with it on and off the proteasome^9,25,43^. How UBE3C retrofits the proteasome and circumvents USP14-mediated suppression to eliminate ultra-stable proteins that otherwise evade proteolysis, and how the seemingly opposing activities of UBE3C and USP14 are integrated on the same proteasome supercomplex, remain completely unresolved.

Here we functionally reconstituted and directly visualized the nonequilibrium functional dynamics of human UBE3C-retrofitted proteasome super-complexes as they degrade an engineered ultra-stable substrate that otherwise evades proteolysis, using time-resolved cryo-EM in combination with biochemical, cellular and mutational analyses. We further captured atomic-level snapshots of highly transient, of substrate-engaged proteasome supercomplexes simultaneously bound to UBE3C and USP14. These studies reveal how UBE3C reprograms the proteasome and counteracts USP14 to enforce the clearance of ultra-stable folds, providing unprecedented insights into the complete functional cycle of human UBE3C-retrofitted proteasome.

## Visualizing UBE3C-retrofitted proteasome

Human UBE3C possesses a basal but relatively weak, Ca^2+^-dependent E3 ligase activity that can ubiquitylate substrates independently of the proteasome^44^. Upon assembly on the proteasome, however, its E4 ubiquitin-elongating activity of UBE3C is markedly enhanced relative to this basal level (Extended Data Fig. 1i). Immunofluorescence imaging of live cells revealed subcellular colocalization of UBE3C, calmodulin and the proteasome (. 1o), and showed that UBE3C presumably formed aggregates in subcellular regions lacking calmodulin. To reconstitute the proteasome-dependent E4 activity of UBE3C, we engineered a model substrate based on a superfolder green fluorescence protein (sfGFP), which is known to resist proteasomal degradation in the absence of UBE3C^20^. Specifically, we fused a tandem repeat of four ubiquitin moieties (Ub4) to the N-terminus of sfGFP and appended an unstructured N-terminal region (residues 1–88) of the yeast Sic1 protein (NtSic1^PY^) to its C-terminus (Extended Data Fig. 1m). To prime this model substrate, designated Ub4-sfGFP-NtSic1^PY^, for E4 activity reconstitution, it was first monoubiquitylated using a ubiquitin mutant with all six lysine residues except Lys48 substituted with alanine (see Methods), thereby ensuring that the subsequent chain extension during UBE3C-catalyzed polyubiquitylation occurs exclusively through Lys48 on the substrate-conjugated monoubiquitin. In our E4 activity assays, the monoubiquitylated model substrate was efficiently polyubiquitylated and then fully degraded by the mixture of UBE3C and the proteasome (Fig. 1a, Extended Data Fig. 1n). By contrast, mutation of the UBE3C active site (C1051A) or omission of UBE3C resulted in deubiquitylation and partial degradation restricted to the NtSic1^PY^ segment, with sfGFP remaining intact, thereby confirming successful reconstitution of UBE3C-dependent enhancement of proteasome processivity (Fig. 1a).

**Figure 1.**
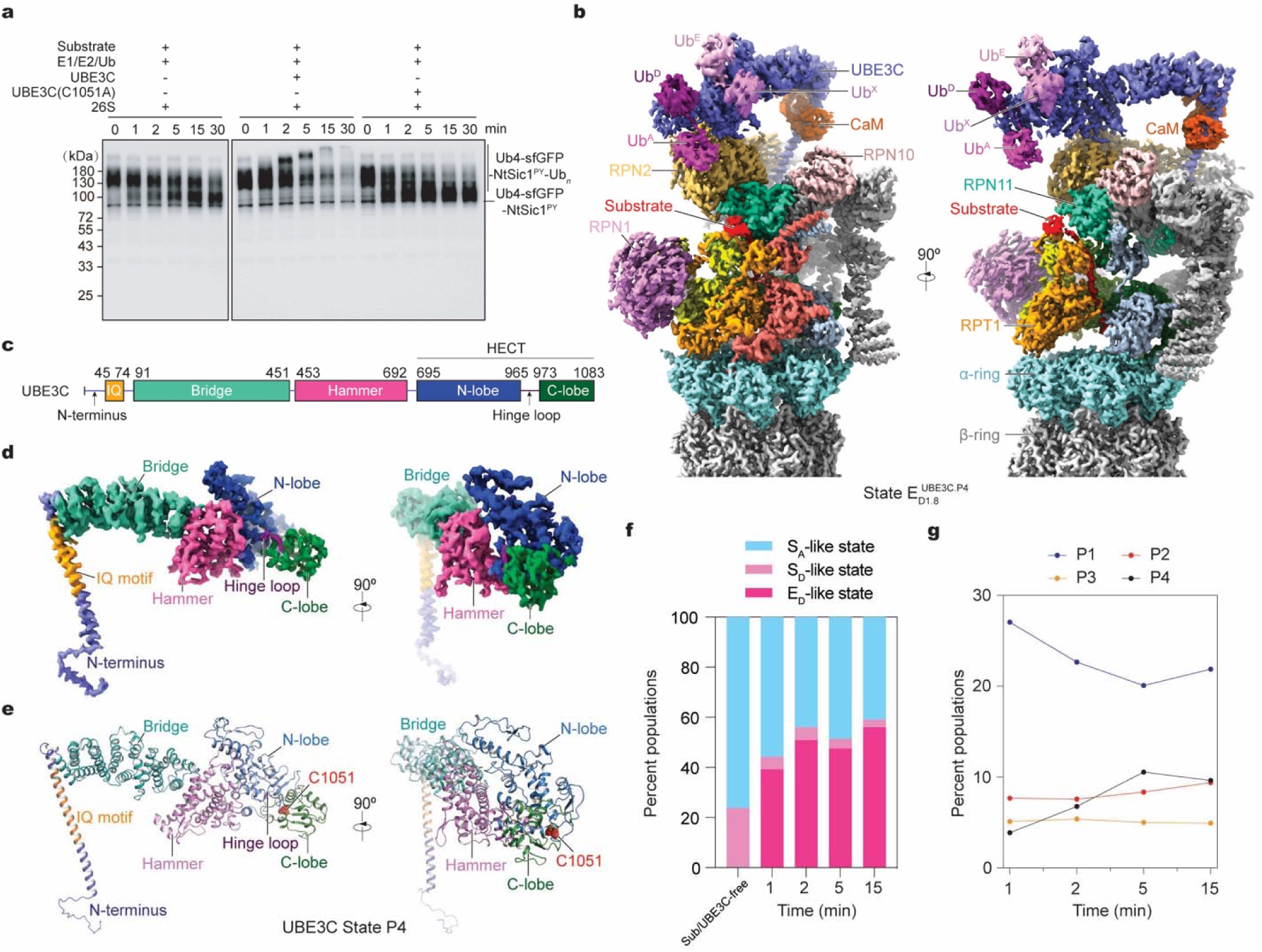
Time-resolved cryo-EM structure of UBE3C-proteasome complex during degradation of a model processivity substrate. **a**, In vitro degradation of a superfolder GFP-based model substrate by human UBE3C-proteasome complex in the presence of E1, E2 and Ubiquitin, analyzed by western blot using an anti-GFP antibody. The substrate, Ub4-sfGFP-NtSic1^PY^-K48Ub*_n_* was generated by partial ubiquitylation of Ub4-sfGFP-NtSic1^PY^ (see Methods). Data are representative of three independent experiments. **b**, Cryo-EM structure of the UBE3C-proteasome complex engaged with the model substrate in state E^UBE3C.P4^. In the right panel, the RPT5 density is omitted to visualize the substrate density inside AAA-ATPase motor. **c**, Domain organization of human UBE3C. CaM, calmodulin; IQ, IQ motif; HECT, Homologous to the E6AP Carboxy Terminus. **d**, **e**, Cryo-EM density map (**d**) and atomic model (**e**) of full-length UBE3C in state P4, extracted from the UBE3C-proteasome complex. **f**, **g**, Kinetic changes in the particle populations of S_A_-like, S_D_-like and E_D_-like states (**f**), and four coexisting UBE3C conformations (**g**), obtained by time-resolved cryo-EM. Particle counts are provided in Extended Data Fig 2.

To visualize substrate-engaged UBE3C-bound proteasome in the absence of USP14, we mixed separately purified human UBE3C and proteasomes with the monoubiquitylated Ub4-sfGFP-NtSic1^PY^ substrate in the presence of E1, E2, and free ubiquitin. The reaction was vitrified 1, 2, 5, and 15 minutes after substrate addition for time-resolved cryo-EM analysis (see Methods). We collected a large cryo-EM dataset of 53,058 micrographs spanning the four time points to enable high-resolution reconstructions. Deep-learning–assisted 3D classification focusing on UBE3C or the proteasomal regulatory particle (RP) resolved four distinct conformational states of UBE3C (P1–P4) at 3.8–4.4 Å resolutions, and 14 conformational states of the proteasome at 2.5–3.6 Å, comprising 3 E_A_-like UBE3C-free, 2 S_D_-like substrate-free, and 9 E_D_-like substrate-engaged conformers (Extended Data Figs. 2-4, Extended Data Table 1). All E_D_-like or S_D_-like proteasome states orthogonally coexist with each of the four UBE3C states, yielding 44 approximately homogeneous conformer subsets with well-defined UBE3C density (Extended Data Fig. 2, and Methods), whereas UBE3C density is largely absent in the E_A_-like states except E^UBE3C^, indicating that UBE3C preferentially recognizes substrate-engaged proteasome conformations (Extended Data Figs. 4a, 5c, d). Six previously uncharacterized E_D_-like proteasome states (E_D1.2_^UBE3C^, E_D1.8_^UBE3C^, E_D1.9_^UBE3C^, E_D2.2_^UBE3C^, E_D3.1_^UBE3C^, and E_D3.9_^UBE3C^) were identified, while three E_D_-like states (E_D0_^UBE3C^, E^UBE3C^, and E^UBE3C^) recapitulated substrate-engaged proteasome conformers observed in the absence of UBE3C^9,10^ (Extended Data Fig. 6a, b). Notably, states E_D4_ and E_D5_ reported for USP14-bound proteasomes were not detected^9^.

### Time-resolved E4-active states of UBE3C

UBE3C has 1083 amino acids, substantially longer than its yeast ortholog Hul5 (910 amino acids), and traverses above the RP, spanning from RPN2 N-terminus to RPN10 and RPN2 C-terminal toroid (Fig. 1b). Its architecture comprises an extended unstructured N-terminal strand that mediates proteasome engagement, a 16-turn long N-terminal helix (H1) harboring the calmodulin-interacting IQ motif, a large arch-shaped Armadillo-like coiled-coil ‘bridge domain’ (BD), a triangular ‘hammer domain’ (HD), and the C-terminal catalytic HECT domain (Fig. 1c-e). The HECT domain consists of an N-terminal lobe (N-lobe) and a C-terminal lobe (C-lobe), with the catalytic Cys1051 residing in the C-lobe. In all four UBE3C conformational states, the N-terminal strand inserts into an interior pocket of the RP, while the HECT N-lobe is clamped between the hammer domain and the HECT C-lobe, with the latter occupying the distal end of the S-shaped UBE3C fold positioned above RPN11 (Figs. 1, 2f-i).

**Figure 2.**
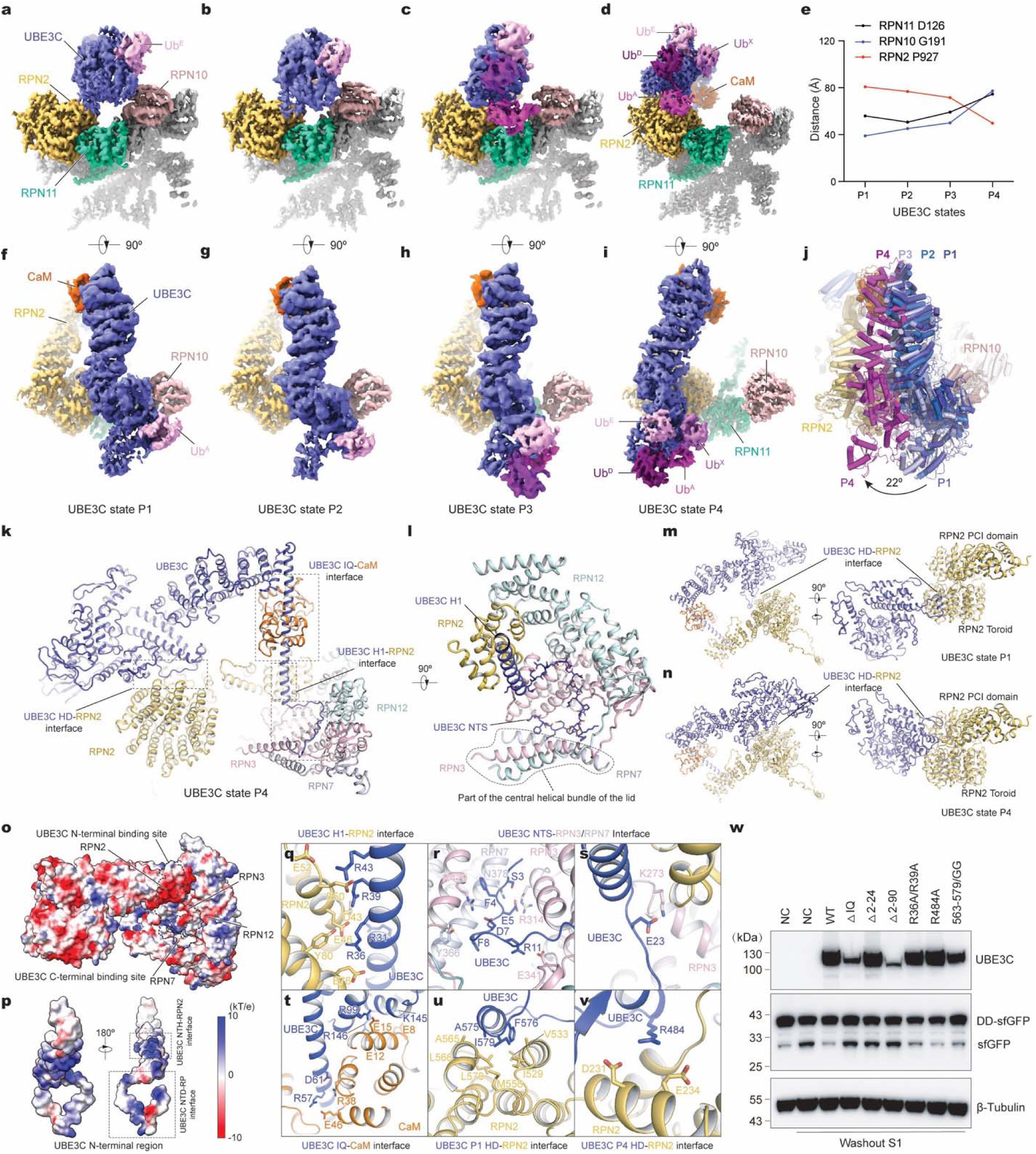
Structure dynamics of proteasome-bound UBE3C in active states. **a**-**d**, Cryo-EM density maps of four distinct UBE3C-lid subcomplexes. **e**, Distances from UBE3C active site to the deubiquitylase RPN11 and the ubiquitin receptor RPN10. Owing to flexibility in the RPN13-RPN2 interface and in the UIM domain of RPN10, densities for these domains were not resolved. Distances were therefore measured from UBE3C active site to Gly192 of RPN10 (nearest to its UIM domain) and to Pro927 of RPN2 (nearest to RPN13). **f**-**i**, Top view of the cryo-EM density maps of four UBE3C states that are rotated by 90° relative to (**a**-**d**). **j**, Structure comparison of proteasome-bound UBE3C in states from P1 to P4, aligned to the proteasome lid and shown from top view. P1 represents the UBE3C conformation positioned closest to RPN10, whereas P4 is positioned closest to RPN2. **k**, Atomic model of UBE3C-calmodulin-RPN2-lid subcomplex in state P4. For clarity, only RPN2 and three lid subunits (RPN3, RPN7 and RPN12) are shown. Dashed boxes indicate the three principal interaction interfaces between UBE3C N-terminal region and the RP subunits. **l,** Closed-up view of UBE3C H1 helix and N-terminal strand inserting into the interior pocket of the lid subcomplex formed between RPN3, RPN7 and RPN12 near the central helical bundle of the lid. **m, n,** Side views of UBE3C-RPN2 subcomplex highlight the interfaces between UBE3C hammer domain (HD) and RPN2 C-terminal domains in state P1 (**m**) and P4 (**n**). **o**, **p**, Electrostatic surface representations of the RPN2 (**o**) and UBE3C N-terminal region (**p**). **q**-**t**, Magnified views of interfaces between the UBE3C H1 helix and RPN2 N-terminal region (**q**), between the UBE3C N-terminal strand (NTS) and RPN3/RPN7 (**r, s**), and between the IQ motif of UBE3C and calmodulin (**t**). All interfaces are shown using the state P4 structure, but their configurations are largely conserved across the four UBE3C states. **u, v**, Structural comparison of the UBE3C hammer domain (HD)-RPN2 interface in states P1 (**k**) and P4 (**l**). **w**, UBE3C enhances proteasomal processing in cells. UBE3C-knockout HEK293T cells stably expressing DD-sfGFP were transiently transfected with UBE3C variants. sfGFP degradation was assessed by western blot using an anti-GFP antibody. Washout S1, the DD ligand Shield removed; NC, negative control (pcDNA3.1); WT, wildtype UBE3C; ΔIQ, IQ motif deletion mutant; Δ2–24 and Δ2–90, N-terminal truncation mutants; 563–579/GG, replacement of residues 563–579 with Gly-Gly. Data are representative of three independent experiments.

Time-dependent quantification of monoubiquitylated Ub4-sfGFP-NtSic1^PY^ substrate processing showed that UBE3C enabled complete degradation of this substrate within approximately 30–60 minutes, with a half-life (*t*_1/2_) of about 5 minutes (Fig. 1a). Consistently, time-resolved cryo-EM revealed that the population of substrate-engaged (E_D_-like) proteasome states rapidly increased to approximately 50% within 2–5 minutes after reaction initiation and remained at 50–60% throughout the observation period (Fig. 1f). During the 2–15-minute interval, the overall occupancy of UBE3C on the proteasome stabilized at around 50%, while its conformational distribution shifted over time (Fig. 1f). Across states P1 to P4, the UBE3C HECT domain progressively migrated away from RPN10 toward RPN2 C-terminal toroid (Fig. 2e). In states P1, P2 and P3, the HECT domain makes brief contact with RPN10. By contrast, in state P4, it was fully detached from RPN10 and docked onto RPN2 above its toroid domain (Fig. 2a-d, m, n). The fractional population of P1 decreased from 28% to 20% between 1 and 5 minutes, while that of P4 increased from 4% to 11% over the same period (Fig. 1g). When UBE3C conformers are aligned against the proteasome subunits and superimposed together, the HECT domain undergoes a rotation of about 22° and exhibits a larger motional amplitude than the N-terminal region of UBE3C (Fig. 2j). Meanwhile, the number of well-resolved ubiquitin moieties bound on the HECT domain increases from one in P1 or P2 to three in P3, and four in P4 (Fig. 2a-i).

Taken together, these observations suggest that P1 represents the initially primed state of proteasome-assembled UBE3C prior to the onset of ubiquitylation. State P2 appears to be an intermediate facilitating the transition from state P1 to P3 or P4. States P3 and P4 likely both correspond to active ubiquitylating configurations compatible with ubiquitin-chain elongation, with P4 accommodating the synthesis of longer ubiquitin chains. Thus, these time-resolved snapshots depict a dynamic functional pathway of UBE3C state transition on the proteasome, in which its conformational reorganization is tightly coupled to progressive ubiquitin loading and processive chain extension.

### Cryptic E3-receptor site

The RP of the proteasome consists of the lid and base subcomplexes^45^. The lid, assembled from nine RPN subunits, adopts a horseshoe-like architecture at the apex of the proteasome, with a central helical bundle formed by the C-terminal helices of all nine RPN subunits at its organizing core^45^. The UBE3C N-terminal strand is almost fully extended and inserts deeply into an interior pocket of the lid near the central helical bundle, featuring three distinct kinks at residues Phe8, Ala19, and Glu24 (Fig. 2k, l). The extreme N-terminal segment (residues 1–24) of UBE3C occupies a narrow cleft between lid subunits RPN3 and RPN7 (Fig. 2k, l). Residues 1–8 pack against the C-terminal helix of RPN7, a component of the central helical bundle of the lid, followed by an approximately 90° turn into a fully extended segment (residues 9–19) that binds an interior pocket of RPN3 and extends toward the PCI domain of RPN12. A second approximately 90° turn then redirects the mainchain toward RPN2, where it transitions into the H1 α-helix of UBE3C (residues 25–82). The N-terminal portion of H1 packs tightly against the N-terminal helical coils of RPN2 (Fig. 2l, q-s, Extended Data Fig. 4e). The interface between the UBE3C N-terminal region (residues 1–50) and RP subunits RPN2, RPN3, RPN7, and RPN12 exhibits electrostatic complementarity and buries a total surface area of 2229 ± 41 Å² that remain invariant across all four UBE3C states (Fig. 2o, p). Notably, the N-terminal strand alone, excluding the H1 helix, already buries 1761 ± 69 Å² on the RP. This interaction is preserved in all E_D_-like and S_D_-like states, as well as in E^UBE3C^, supporting its role as the primary anchoring site for UBE3C recruitment to the proteasome. These observations indicate that UBE3C N-terminal binding site in the lid is already formed in E_A_-like states of the proteasome, but that high-affinity UBE3C association may also depend on its C-terminal interaction with RPN2 or RPN10, as observed in the E_D_-like and S_D_-like states.

In contrast, the interface between the UBE3C hammer domain and the C-terminal toroidal domain of RPN2 is highly variable across different states, burying 583, 717, 676, and 1478 Å² in states P1, P2, P3, and P4, respectively. In state P1, a short helix (residues 575–579) in the hammer domain contacts the T1 site of the RPN2 toroidal domain, which harbors a low-affinity ubiquitin-binding site. In state P4, however, this helix swings away as the entire HECT-hammer assembly shifts upward to contact RPN2 residues Asp231 and Glu234, as well as a loop (residues 269–275) positioned above the T1 site that is disordered in other states (Fig. 2m, n, u, v, Extended Data Fig. 5a, b). Consistent with the highly mobile nature of this interface, mutations in the hammer domain (e.g., R484A or V563G/I579G) did not cause noticeable defects in UBE3C’s E4 activity, whereas mutations in the N-terminal region severely impaired UBE3C’s ability to enforce sfGFP degradation (Fig. 2w; see below).

### Calmodulin regulation of UBE3C

UBE3C contains an IQ motif (residues 45–74) within its N-terminal helix, which serves as a binding site for calmodulin^46,47^. The N-terminal helical region of UBE3C is connected to the bridge domain at the hinge loop (residues 82–91). Right below this loop, the N-terminal lobe of calmodulin wedges into the concave space between the UBE3C N-terminal helix and the bridge domain, structurally supporting UBE3C’s arch-like architecture above the lid (Fig. 2k, t, Extended Date Fig. 8a-l). This intermolecular interaction helps stabilize the overall conformation of UBE3C on the RP. The interface buries 1785, 1726, 1775, and 1816 Å² in states P1, P2, P3, and P4, respectively, which suggests slightly tighter binding in state P4 and supports the role of P4 as the catalytically active state.

Ca^2+^ enhances the E3 ligase activity of isolated UBE3C^44^ (Extended Data Fig. 1h). However, it reduces the E4 activity of UBE3C when associated with the proteasome, presumably by inducing its dissociation from the proteasome complex (Extended Data Fig. 1i-k). To test the functional role of calmodulin binding, we generated UBE3C mutants that affect calmodulin interactions and expressed them in UBE3C-knockout HEK293T cells. These cells stably express a destabilization domain-fused sfGFP (DD-sfGFP) reporter, allowing us to assay proteasome processivity in live cells^20,21,48^ (see Methods). Deletion of the IQ motif (residues 45–74) substantially impaired sfGFP degradation (Fig. 2w, Extended Data Fig. 9f). Likewise, removing N-terminal residues 2–90 (disrupting both the lid and calmodulin interfaces) or residues 2–24 (disrupting only the lid interaction) resulted in the same phenotype—a complete loss of UBE3C-dependent processivity enhancement (Fig. 2w).

### Linkage-specific ubiquitin-binding sites

UBE3C preferentially extends Lys48-linked polyubiquitin chain, while retaining the capacity to assemble Lys29- or Lys33-linked and Lys48/Lys29-branched chains^22,24^. In state P4, four distinct ubiquitin densities are bound to the HECT domain (Fig. 2, 3a-d, Extended Data Fig. 8m-o). Two of these (Ub^A^ and Ub^D^), which are linked via Lys48 of the acceptor ubiquitin (Ub^A^), occupy sites A and D on the C-lobe, respectively (Fig. 3c). The Lys48-linked isopeptide bond lies within approximately 3.5 Å of the catalytic Cys1051 residue, which is well ordered in a P4 reconstruction focused on the ubiquitin densities (Fig. 3j, Extended Data Fig. 4j). This suggests that state P4 captures a snapshot of UBE3C immediately after an isopeptide-bond formation event in which the donor ubiquitin (Ub^D^) has just been transferred to the acceptor (Ub^A^). The other two ubiquitin moieties (Ub^E^ and Ub^X^) bind the N-lobe of the HECT domain at sites E and X, respectively (Fig. 3d). These moieties are putatively linked via Lys29 of the ubiquitin at the X-site, which resides below the E-site. Notably, Lys33 of Ub^X^ is positioned within 3.5 Å of the modelled Lys29-linked isopeptide bond, suggesting that the E/X sites could also accommodate a Lys33-linked diubiquitin in a geometrically compatible configuration (Fig. 3k).

**Figure 3.**
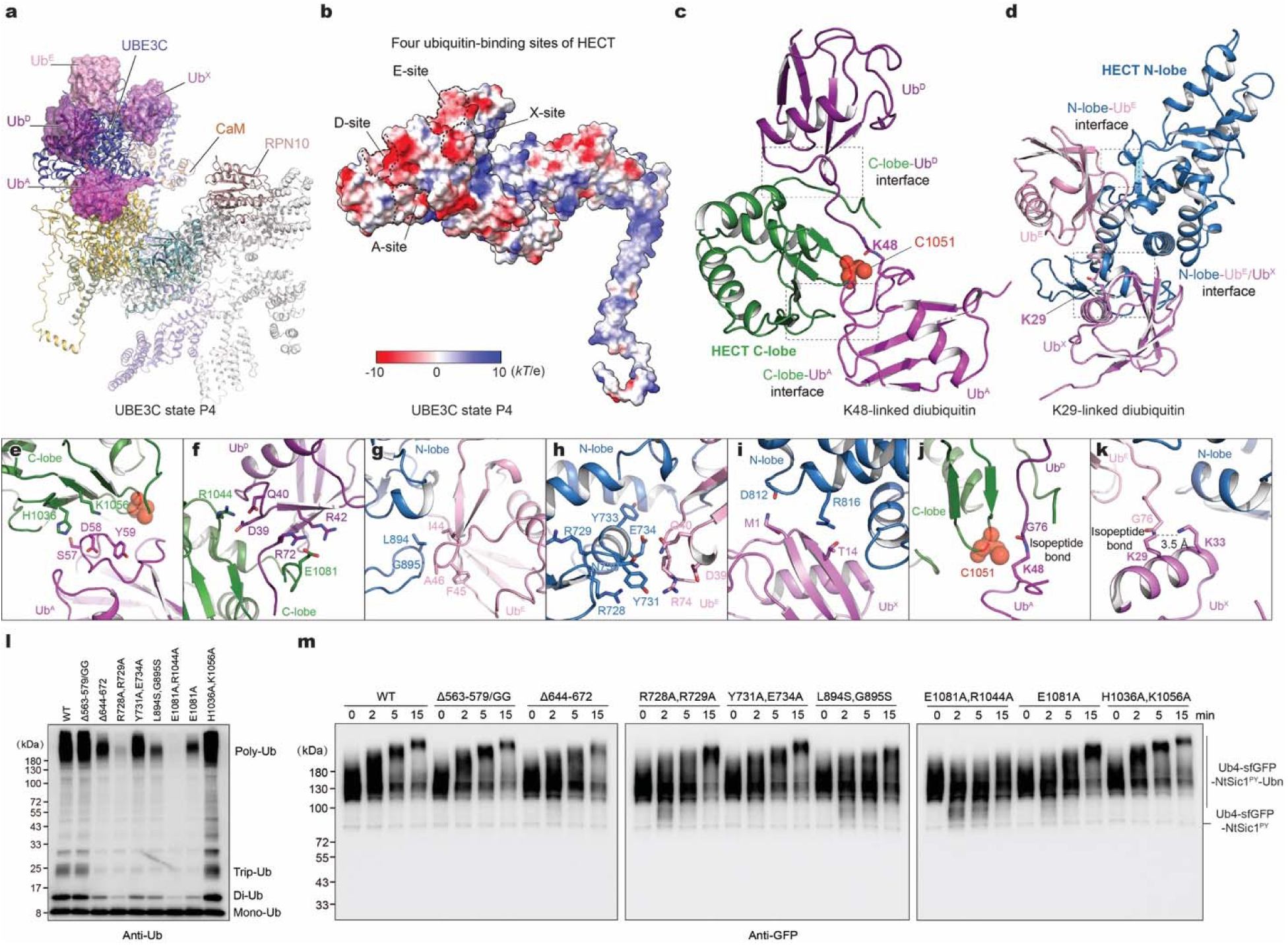
Structural basis of UBE3C-catalyzed ubiquitin-chain elongation and branched chain growth at the proteasome. **a**, Four ubiquitin molecules are highlighted in both cartoon and surface representations in the atomic model of state P4. **b**, Electrostatic surface representation of UBE3C in state P4 showing the acidic or negatively charged surfaces at the four ubiquitin-binding sites designated the A-, D-, E- and X-sites. **c**, **d**, Atomic models of the UBE3C HECT C-lobe with A/D-sites bound to Ub^A^ and Ub^D^ linked by K48 isopeptide bond (**b**) and the UBE3C N-lobe with E/X-sites bound to Ub^E^ and Ub^X^ linked by K29 isopeptide bond (**c**). The UBE3C active site residue Cys1051 is shown as a sphere. **e-i**, Closed-up views of the C-lobe-Ub^A^ interface (**e**), the C-lobe-Ub^D^ interface (**f**), the N-lobe-Ub^E^ interface (**g**, **h**), and the N-lobe-Ub^X^ interface (**i**). **j**, **k**, Closed-up views showing the geometry of K48- and K29-linked isopeptide bond models at the UBE3C C-lobe-Ub^A^ interface (**d**) and UBE3C N-lobe-Ub^X^ interface (**e**), respectively. **l,** In vitro ubiquitylation activity of UBE3C mutants. Reactions were analyzed by SDS-PAGE and immunoblotting with an anti-ubiquitin antibody. Experiments were repeated three times with consistent results. **m**, Ubiquitylation and degradation of Ub4-sfGFP-NtSic1^PY^-K48Ub*_n_* by the proteasome in the presence of UBE3C mutants. Reactions were analyzed by SDS-PAGE and immunoblotting with an anti-GFP antibody. Data are representative of three independent experiments.

Remarkably, in state P3, the A-site of the HECT domain is positioned approximately 55–60 Å above the ubiquitin-binding site of RPN11—a distance comparable to the dimension of a single ubiquitin moiety (Fig. 2c, Extended Data Fig. 8q). This geometric arrangement implies that an elongated ubiquitin chain bound at the A-site could be readily handed off to RPN11, thereby facilitating substrate delivery to the AAA-ATPase translocation pathway for processive unfolding. Moreover, low-pass filtering of the P3 or P4 cryo-EM maps reveals low-resolution density resembling a forked, tree-like appearance, with one branch extending toward the A/D sites on the C-lobe, and the other extending toward the E/X sites on the N-lobe (Extended Data Fig. 8p, q). This configuration suggests that states P3 and P4 are compatible with a “ubiquitin-chain branching” process, in which the catalytic C-lobe of the HECT domain pivots between a down conformation that supports K48-linked chain synthesis at the A/D sites and an up conformation that would enable K29-linked chain synthesis at the E/X sites. Although we did not directly observe the C-lobe up conformation in our cryo-EM reconstructions, structural studies of other HECT-type ligases have visualized an analogous C-lobe up conformation that is geometrically compatible with this model (Extended Data Fig. 8s-w)^49,50^.

The ubiquitin–UBE3C interfaces at sites A, D, E, and X bury surface areas of 674, 690, 1173, and 337 Å², respectively. In line with canonical ubiquitin–receptor interactions^51,52^, these ubiquitin-binding sites are predominantly acidic, negatively charged, complementing the electrostatic surfaces of their ubiquitin counterparts (Fig. 3b). The ubiquitin bound at the E-site—previously identified as an exosite in other HECT-type ligases^52–54^—engages UBE3C through two distinct interfaces: one in which N-lobe residue Leu894 interacts with the Ile44 hydrophobic patch of ubiquitin, and a second involving residues Glu734 and Tyr733 (Fig. 3g, h). Ubiquitin density at the E-site is observed across all four UBE3C conformational states, whereas Ub^A^ and Ub^D^ densities are well-resolved in state P3 as well as P4, and Ub^X^ is clearly discernible only in P4 (Fig. 2a-i).

To examine the functional importance of these ubiquitin-binding sites, we performed site-directed mutagenesis at the UBE3C–ubiquitin interface. A double mutant (E1081A/R1044A) at the D-site on the C-lobe resulted in a severe loss of ligase activity (Fig. 3l, m). Mutations at the E-site (R728A/R729A, Y731A/E734A, and L894S/G895S) also reduced the ligase activity, albeit less dramatically than the D-site mutant. To investigate linkage specificity, we analyzed *in vitro* ubiquitylation products using linkage-specific antibodies. Notably, several E-site mutations (Y731A/E734A and L894S/G895S) disrupted K29- and K33-linked ubiquitylation much more strongly than K48-linked synthesis (Extended Data Fig. 9h). These data support the hypothesis that the E/X sites are specialized for K29/K33-linked chain formation. Collectively, our structural and functional analyses demonstrate that the four ubiquitin-binding sites on the HECT domain are specialized to promote ubiquitin-chain extension with distinct linkages and, when acting together, enable the assembly of branched ubiquitin chains with hybrid linkages.

### Visualizing transient UBE3C-USP14-26S supercomplex

A previous study has found that UBE3C and USP14 are prone to co-cycle on and off the proteasome during substrate processing^43^, complicating efforts to reconstitute a stable UBE3C-USP14-26S super-complex. To overcome this hurdle, we engineered another model substrate in which the N-terminal segment of Sic1^PY^ is fused to a mutated sfGFP variant, NtSic1^PY^-cp8sGFP, that contains cp8 mutations in GFP (see Methods). In contrast to the degradation-resistant sfGFP, NtSic1^PY^-cp8sGFP was more readily degraded by the proteasome even without UBE3C (Extended Data Fig. 11j). When both UBE3C and USP14 are present, UBE3C counteracts USP14-mediated suppression and restores proteasomal degradation of NtSic1^PY^-cp8sGFP (Extended Data Fig. 11j, k).

To capture highly transient UBE3C-USP14-26S supercomplexes, we incubated human proteasomes with separately purified UBE3C and USP14 in a reaction buffer containing E1, E2, ubiquitin and polyubiquitylated NtSic1^PY^-cp8sGFP. The mixtures were then vitrified 15, 30, 60, 120 and 300 seconds after substrate addition for time-resolved cryo-EM analysis (Extended Data Fig. 11l). We collected a considerably large single-particle dataset of micrographs from these cryo-EM samples that, after exhaustive focused 3D classification, yielded 2 UBE3C states, 3 USP14 states, and 10 proteasome states (Fig. 5a-h, Extended Data Figs. 11a, 12).

Notably, USP14 occupancy on the proteasome decreased sharply within the first 5 minutes of degradation, whereas the dissociation of UBE3C was much slower than USP14 (Extended Data Fig. 11l). Likely owing to the greater conformational dynamics of the substrate-engaged UBE3C-USP14-26S supercomplex and the relatively low UBE3C occupancy on the proteasome, the 10 proteasome states reconstructed here correspond to a subset of the proteasomal conformers as observed in the UBE3C-26S supercomplex. Except for states E_D1.2_, E_D1.8_ and E_D1.9_, all proteasome states observed for UBE3C-26S without USP14 were also observed herein, whereas UBE3C were also observed to sample states P1 and P3 that were convoluted with those proteasome states (Extended Data Figs. 11, 12).

### UBE3C bypasses USP14 for ubiquitin shuttling

Despite adding USP14 at stoichiometric saturation in our time-resolved cryo-EM analysis, in the proteasome state E_D2.1_, only a small fraction (14.5%) of proteasomes display stable cryo-EM density for the USP domain of USP14 bound to the OB ring of the AAA-ATPase motor (designated E_D2.1_^UBE3C-USP14-Ub^), whereas the ubiquitin-binding pocket of the USP domain appears to be empty in 9.7% of the population (designated E_D2.1_^UBE3C-USP14^, see Fig. 4a, b, e, f). The majority (75.8%) of E particles, designated E_D2.1_^UBE3C-UBL^, lack discernible cryo-EM densities for the entire USP domain, with only the UBL domain of USP14 remaining visible on RPN1 (Fig. 4c, g).

**Figure 4.**
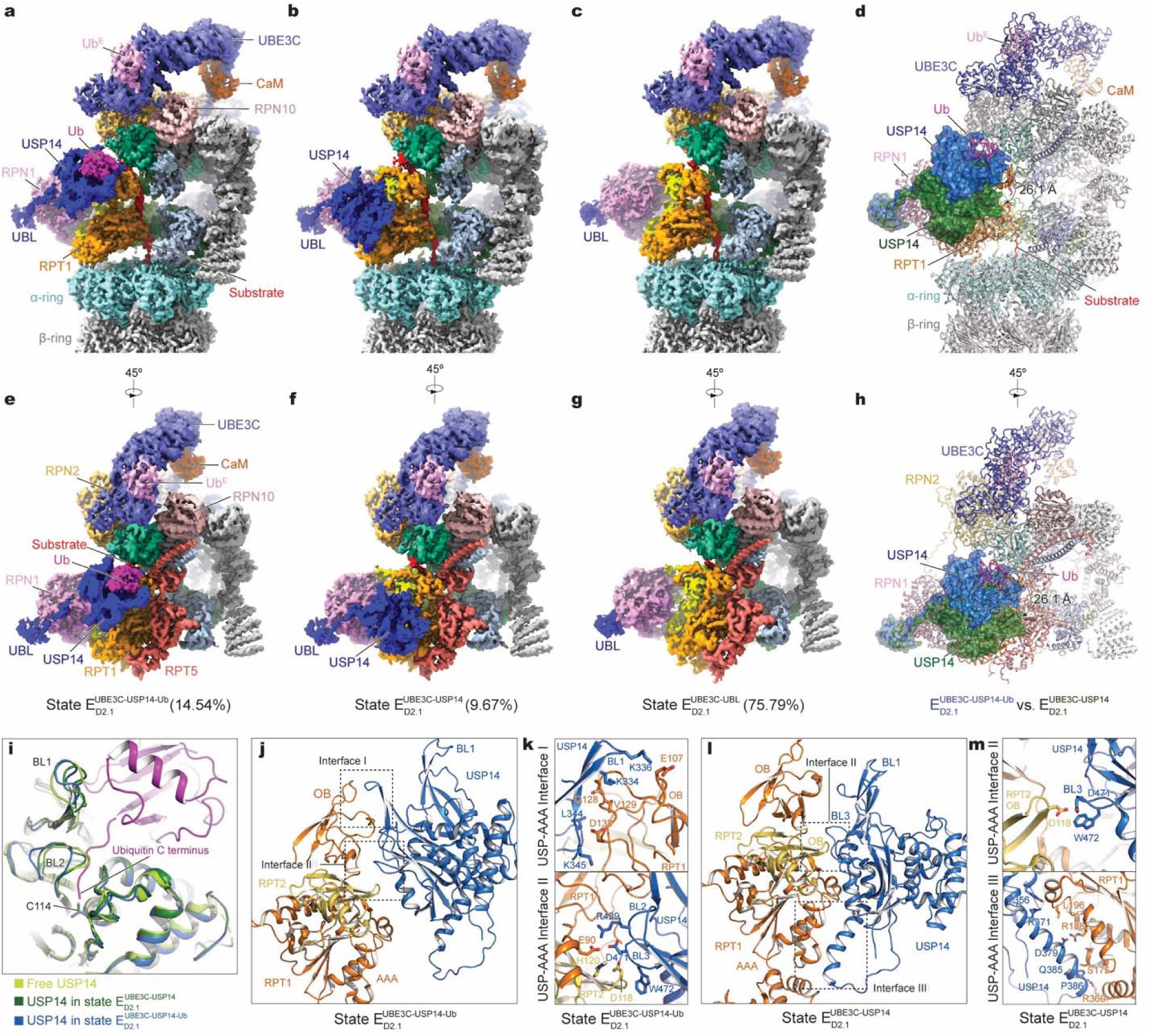
**Functional interplay of USP14 and UBE3C by time-resolved cryo-EM. a-c**, Composite cryo-EM density maps of three distinct UBE3C-USP14-26S complexes in the proteasome state E_D2.1_. The proteasome and USP14 densities correspond to focused 3D subclasses of the state E_D2.1_ class, whereas the UBE3C density corresponds to the UBE3C P1 state refined from focused 3D subclasses from all E_D_/S_D_-like particles (see Methods). **d,** Structural comparison of proteasome-bound USP14 in E_D2.1_ states, aligned to the proteasome lid. USP14 is highlighted in both cartoon and surface representations. **e-g,** Rotated views of the cryo-EM density maps shown in **a-c**. **h,** Rotated view of the atomic model that shown in **d**. **i,** Structural comparison of the blocking loops of USP14, obtained by superimposing the USP domains in the E_D2.1_ states with the crystal structure of free USP14 (PDB ID: 2AYN). **j-m,** Side by side comparison of USP14 interactions with RPT1-RPT2 between states E_D2.1_^UBE3C-USP14-Ub^ (ubiquitin-bound USP14 state; panel **j**, **k**) and E_D2.1_^UBE3C-USP14^ (ubiquitin-free USP14 state; panel **l**, **m**). The latter shows that the BL1 of USP14 has dissociated from the OB domain of RPT1, where they are in close contact in the ubiquitin-bound state E_D2.1_^UBE3C-USP14-Ub^.

**Figure 5.**
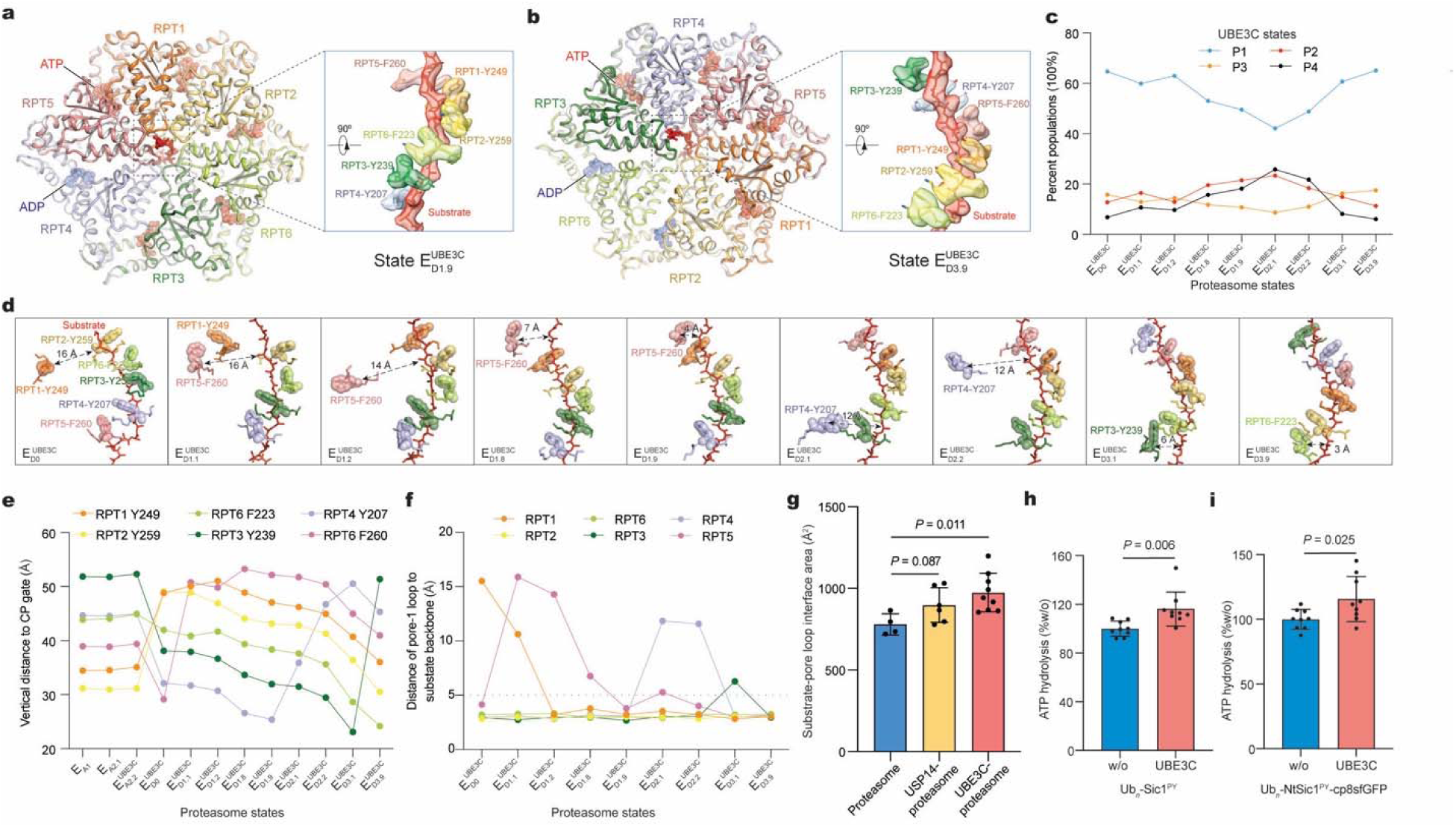
S**t**ructural **dynamics of allosteric regulation of the AAA-ATPase motor by UBE3C**. **a, b**, Atomic models of the AAA-ATPase motor in states ED1.9^UBE3C^ and ED3.9^UBE3C^ in cartoon representation. Right insets show the zoomed-in side views of pore-1 loops contacting the substrate backbone, with both cryo-EM densities (transparent surface representation) and atomic models (stick representation) displayed. **c**, Percent populations of proteasome-bound UBE3C states across distinct proteasome-UBE3C complex conformations. Intersection data of UBE3C and proteasome classifications were used. **d**, Architecture of the pore-1 loop staircase engaging the substrate in distinct proteasome-UBE3C complex states. The longest pore-1 loop-substrate distance is indicated for each state. Pore-1 loop residues are shown in stick representation, with aromatic residues additionally highlighted as transparent spheres. **e**, **f**, Distances from the pore-1 loop of each ATPase subunit to the CP gate (**e**) and to the substrate (**f**) in different states of UBE3C-proteasome complex. **g,** The interfacial areas buried between the substrate backbone and the pore loops of ATPases in all E_D_-like states of UBE3C-bound proteasome in comparison with the previously reported two E_D_-like proteasome states without UBE3C. **h, i**, ATPase activity of proteasome during degradation of Ub*_n_*-Sic1^PY^ and Ub*_n_*-NtSic1^PY^-cp8sfGFP. *P* values were calculated relative to w/o (without UBE3C) using a two-tailed unpaired *t*-test. Data are mean ± s.d. from three independent experiments, each performed in triplicate.

Ubiquitin-binding to the USP domain, which sandwiches its blocking loops 1, 2 and 3 (BL1–3) with the OB ring of the AAA-ATPase, is critical for stabilizing USP14 association with the proteasome^9^. In state E_D2.1_^UBE3C-USP14^, where the USP domain is devoid of ubiquitin, BL1 and BL3 are largely dissociated from the OB domains of RPT1-RPT2 (Interface I; Fig. 4i-l), while BL3 retains only a minimal contact with the OB domain of RPT2 (Interface II, Fig. 4k, m). Notably, the ubiquitin-free USP domain in E_D2.1_^UBE3C-USP14^ is displaced by 26.1 Å away from the OB ring toward the AAA domain of RTP1 relative to the ubiquitin-bound USP in E_D2.1_UBE3C-USP14-Ub (Fig.4d, h), thereby allowing a larger interfacial contact formed between the RPT1 AAA and USP domains (Interface III; Fig. 4m). The ubiquitin-free USP14-RPT1-RPT2 conformation in E_D2.1_^UBE3C-USP14^ is markedly distinct from that of the USP14-bound proteasome in the absence of UBE3C^9^.

The binding mode of UBE3C at the proteasome positions the catalytic C-lobe of its HECT domain directly above the intrinsic DUB RPN11, placing their ubiquitylating active site within a length of roughly one to two ubiquitin moieties away from the deubiquitylating active site of RPN11. By contrast, the deubiquitylating site of USP14 in the proteasome lies more than four ubiquitin lengths away, posing a structural disadvantage for USP14 to intercept the ubiquitin transfer from either UBE3C or the nearby ubiquitin receptors RPN10/RPN13 en route to RPN11. Because the HECT C-lobe of UBE3C is positioned at approximately the same latitude as the RPN10 UIMs and RPN13 on RPN2 and shuttle between RPN10 and RPN2, it can presumably hand off newly elongated polyubiquitin chains to the ubiquitin receptors and RPN11. These geometric constraints leave USP14 with little opportunity to engage the substrate-proximal ubiquitin and to divert the productive substrate-translocation pathway, when ubiquitin chains are newly synthesized on substrate by proteasome-bound UBE3C. Taken together, our structural and kinetic data indicate that UBE3C association with the proteasome promotes USP14 recycling by minimizing the probability of ubiquitin transfer to USP14 and overriding its ubiquitin-binding priority, thereby suppressing the DUB activity of USP14 while retaining UBE3C-mediated control over the proteasome.

### UBE3C promotes AAA unfoldase activity

Several newly identified E_D_-like states of the AAA-ATPase motor —E_D1.8_, E_D1.9_, E_D3.9_ and E_D3.1_—consistently exhibit tighter substrate–pore loop interactions than previously characterized conformers^9,10^ (Fig. 5a, b, d-f). States E_D3.9_ and E_D3.1_ were observed both in the absence and presence of USP14, whereas E_D1.8_ and E_D1.9_ appeared exclusively in the UBE3C-26S dataset in the absence of USP14. States E_D1.9_ and E_D3.9_ represents the most extreme cases, with all six pore-1 loops from the ATPase subunits simultaneously contacting the substrate (Fig. 5a, b). In state E_D1.9_, all six pore-1 loops form a spiral staircase from RPT5 (top) to RPT4 (bottom), whereas state E_D3.9_ shows a pore-loop spiral from RPT3 (top) to RPT6 (bottom). In these fully engaged states, Van der Waals contacts between pore loops and the substrate backbone are estimated to increase by 30–50% compared with any previously reported UBE3C-free states in which only four or five subunits contact the substrate^9,10^ (Fig. 5g). This strengthened grip allows the ATPase motor to exert greater mechanical forces, sufficient to unfold an ultra-stable protein such as sfGFP. Consistent with this higher-affinity engagement, all six ATPase subunits in these states display well-defined cryo-EM densities for nucleotides, with the lowest subunits in the spiral bound to ADP (Extended Data Fig. 7). By contrast, all previous substrate-engaged states reconstructed without UBE3C, with or without USP14, have consistently showed one or two ATPase subunits disengaged from substrates and in apo-like or poorly ordered nucleotide states^9,10^. These findings suggest that the fully engaged intermediates are either transiently stabilized or allosterically induced by UBE3C.

To quantify the extent to which UBE3C remotely modulates the AAA-ATPase motor via long-range allostery, we measured the ATPase activity for UBE3C-loaded proteasomes during degradation of polyubiquitylated substrates, such as Sic1^PY^ and NtSic1^PY^-cp8sfGFP, in the absence of E1, E2, and free ubiquitin (see Methods). The presence of UBE3C alone, without involving its E4 chain-elongation activity, enhanced proteasomal ATPase activity by approximately 10-20% (Fig. 5h, i), confirming that UBE3C imposes a non-catalytic, long-range allosteric regulation on the AAA-ATPase motor.

The HECT domain of UBE3C swings between RPN2 and RPN10 above RPN11. Because all four UBE3C conformational states were represented within each E_D_-like proteasome state, we wondered whether specific UBE3C states preferentially associate with particular proteasome conformers. We therefore quantified the populations of the four UBE3C conformers across the nine E_D_-like proteasome states (Fig. 5c). UBE3C states P1 and P3 reach their lowest abundance in state E_D2.1_ compared with the other proteasome states, whereas states P2 and P4 are maximally populated in proteasome state E_D2.1_. The populational changes of UBE3C states also gradually increase or decrease in line with the presumed order of AAA-ATPase motor state transitions, suggesting that UBE3C conformational changes are coupled to the ATPase cycle in the proteasome.

### Insights into UBE3C-reprogrammed proteasome

Our comprehensive data reveal how UBE3C-retrofitted proteasome is reprogrammed at thermodynamical, kinetic, and allosteric levels (Fig. 6a). Thermodynamically, in situ synthesis of ubiquitin chains and branched chain topologies by UBE3C at the proteasome promotes multivalent interactions of ubiquitin moieties with proteasomal ubiquitin receptors, markedly increasing substrate-binding affinity on the proteasome and preventing premature substrate release before degradation is accomplished (Fig. 6b).

**Figure 6.**
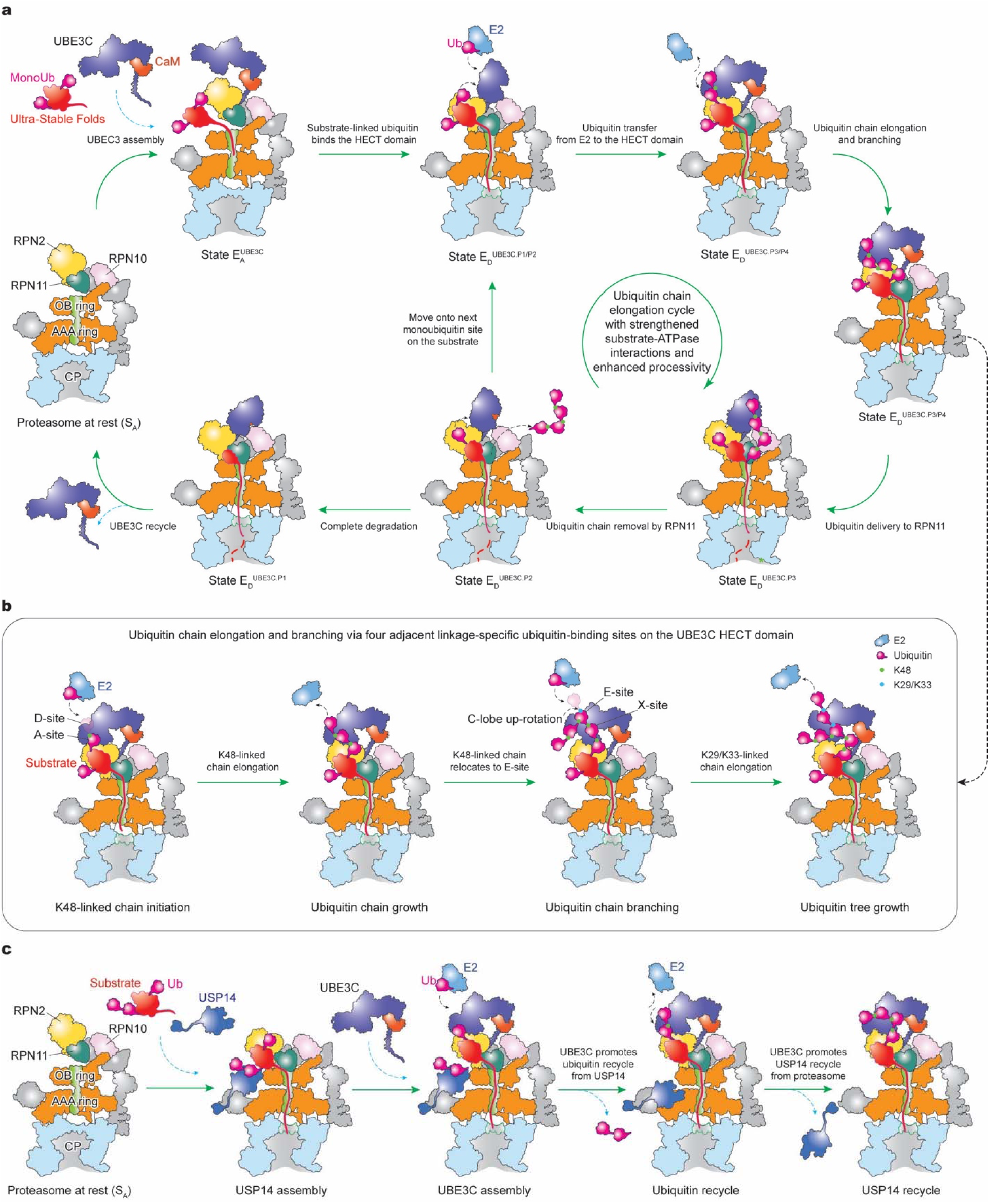
Schematics showing the proposed model of complete functional cycle of UBE3C-retrofitted proteasome. a, Monoubiquitylated substrate can engage with the proteasome with or without the presence of UBE3C. However, calmodulin-regulated UBE3C assembly on the proteasome that already captured a substrate prevents a premature release of the substrate, when the substrate-linked monoubiquitin is captured by the proteasome-associated UBE3C via four ubiquitin-binding sites on its catalytic HECT domain. Once primed and fully ready, UBE3C catalyzes transfer of ubiquitin from E2 enzyme to the substrate, growing a ubiquitin chain or building a branched chain starting on the monoubiquitin. Given that the catalytic C-lobe of UBE3C HECT domain positioning right above the deubiquitylase RPN11 with a distance of one to two ubiquitin moieties, and between ubiquitin receptors RPN10 and RPN13, the newly synthesized ubiquitin chains can be quickly captured by nearly ubiquitin receptors and be delivered to RPN11, with the minimal diffusion distance for enhanced kinetics for ubiquitin delivery. It is also possible that the substrate-conjugated ubiquitin delivery to RPN11 can bypass the ubiquitin receptors for even faster kinetics. RPN11 subsequently catalyzes removal of ubiquitin chain for processive substrate unfolding and degradation. Once the mission is accomplished, UBE3C can be recycled and released from the proteasome via calmodulin and calcium-dependent regulation. **b**, A hypothetical model for UBE3C-catalyzed ubiquitin chain elongation and tree growth via four adjacent ubiquitin-binding sites on the UBE3C HECT domain. Given that the A-site on the HECT C-lobe is the closest to the substrate-translocation entrance of the AAA-ATPase motor in the proteasome, ubiquitin binding to the A-site occurs at the same time of substrate-engagement with the AAA-ATPase motor, which is preferably used by UBE3C to synthesize K48-linked isopeptide bond for chain elongation, with E2-transferred ubiquitin binding the D-site before isopeptide bond formation. When the chain is long enough, some ubiquitin moiety in the chain can diffuse to and navigate the nearby E/X-sites on the HECT N-lobe, which can be added with E2-transferred ubiquitin via K29 or K33-linked isopeptide bond, thus creating a branched, tree-like chain. **c**, A schematic model of USP14 promoting UBE3C assembly on the proteasome and then recycling from the proteasome due to the effect that UBE3C reduces the chances of ubiquitin delivery to USP14.

Kinetically, UBE3C accelerates proteasomal degradation by aligning its catalytic C-lobe above RPN11 and between the ubiquitin receptors RPN10 and RPN13 in a quaternary arrangement that co-localizes ubiquitylation and deubiquitylation activities. This architecture allows newly assembled ubiquitin chains to be transferred to RPN11 over an extreme shortcut of diffusion path—on the order of one to two ubiquitin lengths—thereby bypassing USP14 altogether and maximizing catalytic efficiency.

Allosterically, UBE3C binding enhances AAA unfoldase activity and boosts the substrate-unfolding forces of the proteasome by 30–50%, allowing the ATPase motor to ultimately surmount the relatively high unfolding energy barriers of ultra-stable protein folds. UBE3C preferentially associates with substrate engaged E_D_ like proteasome states, with its occupancy in E_D_-like proteasome states nearly tenfold higher than in E_A_-like states (Extended Data Fig. 5c). The proteasomal lid serves as an E3 receptor platform that physically separates local UBE3C motions from the AAA-ATPase motor, creating an allosteric relay that establishes a positive feedback loop between UBE3C-catalysed ubiquitylation with ATPase-driven substrate unfolding. This allosteric configuration biases UBE3C toward the substrate-engaged, interaction-strengthened states of the AAA-ATPase motor that suppress backward slippage during substrate translocation, sustaining heightened processivity while shortening idle intervals between catalytic bursts.

Although USP14 association appears to promote UBE3C loading onto the proteasome, UBE3C in turn overrides the ubiquitin-binding priority of USP14, thereby facilitating USP14 recycling off the proteasome (Fig. 6c). This allosteric antagonism grants UBE3C the capability to expel USP14 from the proteasome and effectively invert the direction of proteasome regulation. Once degradation is complete, UBE3C can dissociate from the proteasome through a Ca^2+^ dependent mechanism^44^. Calcium binding remodels calmodulin conformation and weakens its interaction with the IQ motif in the UBE3C N-terminal helix^55^, increasing the conformational entropy of UBE3C and triggering its release from the proteasome. Thus, UBE3C-mediated potentiation of proteasome function operates within a regulatory framework that specifically enforces proteolysis of ultra-stable folds that pose proteotoxic and pathological threats to cells, while calmodulin—which binds UBE3C but not the proteasome—provides a critical Ca^2+^ sensitive regulatory layer governing the coupled dissociation of UBE3C and USP14 from the proteasome.

By coupling intracellular Ca^2+^ signaling to UBE3C recycling from the proteasome, cells can dynamically tune this conserved ubiquitylation–degradation-coupled pathway to match physiological demands. Elevated intracellular Ca^2+^ and early protein aggregation are recurrent hallmarks of neurodegenerative disorders^56,57^, yet the molecular connections between these phenomena have remained elusive. Our findings support a Ca^2+^–UBE3C–proteasome regulatory axis in which Ca^2+^ elevation promotes premature UBE3C dissociation, thereby attenuating the proteasomal clearance of early aggregation-prone substrates and linking ionic dysregulation to proteostasis failure.

Together, these exquisite integrated mechanisms clarify how the highly dynamic UBE3C-proteasome interactions reprogram the UPS into in a globally upregulated, highly efficient proteolytic holoenzyme machine, thereby boosting degradative capacity to overcome ultra-stable folds. Consequently, UBE3C-reprogrammed proteasome is particularly advantageous and indispensable under proteotoxic stress, when rapid clearance of aggregation-prone substrates is critical. Our analysis further revealed for the first time that UBE3C utilizes four closely apposed, linkage-specific ubiquitin-binding sites to drive both ubiquitin-chain extension and branching (Fig. 6b). Given the conservation of the HECT domain structure, the structural basis of branched chain synthesis elucidated here for UBE3C potentially represents a generic paradigm that could be similarly exploited by other E3 ligases to assemble hybrid branched ubiquitin chains.

The capacity of UBE3C to enforce degradation of ultra-stable proteins like sfGFP has far-reaching implications in health and disease. The core structure of sfGFP adopts a β-barrel fold, and many proteolysis-resistant aggregates in neurodegenerative disorders likewise form extensive, densely packed β-sheet architectures^7,8^. Thus, sfGFP serves as an informative model for how UBE3C-retrofitted proteasomes may engage and dismantle such ultra-stable misfolded species in vivo. Dysregulation of UBE3C in neurodegeneration may contribute to pathological progression^12,13^, whereas UBE3C overexpression in cancer may enable tumor cells to cope with elevated proteostasis demands or evade apoptosis^15,24^. Restoring UBE3C activity in neurodegenerative disease and inhibiting UBE3C in cancer are therefore both promising therapeutic strategies. We anticipate that the atomic structures and functional dynamics of UBE3C–26S and UBE3C-USP14-26S super-complexes described here will facilitate the development of proteolysis-targeting agents that harness this mechanistically coupled E3–DUB-proteasome axis as an innovative therapeutic platform.

## Methods

### Constructs

The full-length human *UBE3C* coding sequence was cloned into a modified pGenLenti vector containing an N-terminal 6×His tag followed by a Tobacco etch virus (TEV) protease cleavage site. The resulting construct was used for lentiviral packaging and generation of a stable HEK293T cell line. For transient expression and protein purification, *UBE3C* and its mutants were subcloned into pcDNA3.1 with an N-terminal 6×His–biotin tag and TEV protease site (HBT). Full-length human *calmodulin* was cloned into pcDNA3.1. The recombinant proteasome substrates were based on sfGFP reporters fused to defined degron elements.

Ub4-sfGFP-NtSic1^PY^ comprises an N-terminal linear tetra-ubiquitin chain, superfolder GFP (sfGFP)^58^, and a C-terminal intrinsically disordered region derived from Saccharomyces cerevisiae Sic1 (amino acids 1–88; NtSic1), followed by a T7 tag and a 6×His tag. In the Ubl-sfGFP-NtSic1^PY^, the tetra-ubiquitin chain was replaced with the ubiquitin-like (UBL) domain of human RAD23B (amino acids 1–140). NtSic1^PY^-cp8sGFP consists of NtSic1 fused at its C-terminus to cp8sGFP, a circularly permuted GFP variant that is well folded but thermodynamically less stable than sfGFP^59^. The PY motif (Pro-Pro-Pro-Ser) appended to the C-terminus of NtSic1 enables recruitment of the WW-HECT E3 ubiquitin ligase (derived from S. cerevisiae Rsp5) to catalyze substrate polyubiquitylation^60^. All bacterial substrates were codon-optimized for expression in Escherichia coli and cloned into the pET-28a vector. The recombinant substrate DD-sfGFP, which contains an N-terminal destabilizing domain (DD) conferring small molecule ligand-dependent stability derived from the L106P mutant of human FK506-binding protein 1a (FKBP1A), and a C-terminal sfGFP, was synthesized with codon optimization for expression in human cells and then cloned into pcDNA3.1/Hygro vector. All constructs used in this paper were synthesized and cloned by GenScript.

### Cell culture

The stable HEK293 cell line expressing HTBH (6×His, TEV protease site, Biotin and 6×His)-tagged RPN11 was a gift from L. Huang as previously generated^61^. The Expi293 cell line and Hela cell line were gifts from the laboratory of J. Sodroski and C. Li, respectively.

HEK293T cell line was obtained from BMCR (National BioMedical Cell Resource Center). All cell lines were confirmed to be free of mycoplasma contamination. The table HEK293 cells, HEK293T cells and Hela cells were cultured in DMEM (Gibco) supplemented with 10% fetal bovine serum (Gibco) and 1% penicillin/streptomycin at 37 °C in 5% CO_2_. Expi293 cells were cultured in SMM 293-TII medium (SinoBiological) at 37 °C in 5% CO_2_.

### Stable cell line generation

To generate the DD-sfGFP HEK293T cell line, HEK293T cells were transiently transfected with pcDNA3.1/Hygro-DD-sfGFP plasmid using Universal Transfection Reagent (Yeasen). After 48 h, the cells were cultured in medium containing 100 µg ml^-1^ hygromycin for an additional 20 d to select for cells that had undergone successful integration. GFP-positive cells isolated through cell sorting using the BD Aria Fusion flow cytometer and subsequently used establish a UBE3C-knockout (KO) cell line employing EZ-Editor^TM^ Kit (Ubigene) according to the manufacturer’s protocols. For the generation of the HTBH-RPN11 Expi293 cell line, Expi293 cells were co-transfected with pQCXIP-HTBH-RPN11 (a gift from D. Zhang), along with psPAX2 and pMD2.G. Viruses were harvested and used to infect Expi293 cells.

### Protein Expression and purification

Human proteasome was purified from stable HEK293 or Expi293 cell line with HTBH-tagged RPN11 as previously described^9,10,45,62^. Briefly, cells were harvested and homogenized in a homogenization buffer (50 mM PBS (77.4% Na_2_HPO_4_, 22.6% NaH_2_PO_4_, pH 7.4), 0.5% NP-40, 1 mM DTT, 5 mM ATP, 5 mM MgCl_2_, 10% glycerol and protease inhibitor cocktail (Roche).

After centrifugation, the supernatant was mixed with Streptavidin affinity resin (Yeasen) at 4 °C for 3 h. The resin was loaded to a gravity column and washed with 20 column volumes (CV) of homogenization buffer, followed by 10 bed volumes of TEV cleavage buffer (50 mM Tris-HCl [pH 7.5], 5 mM ATP and 5 mM MgCl_2_). Then proteasomes were cleaved from the beads by TEV protease (Invitrogen) and further purified by SEC (size-exclusion chromatography) using a Superose 6 10/300 GL (GE Healthcare) in FPLC buffer (30 mM HEPES [pH 7.5], 60 mM NaCl, 0.5 mM DTT, 1mM MgCl_2_, 0.6mM ATP, 10% glycerol).

For Extended Data Fig. 1a-c, UBE3C was purified from a stable HEK293T cell line with 6×His and TEV tagged UBE3C. Cells were harvested at 80-90% confluency and lysed by Dounce-homogenizing in lysis buffer (50 mM PBS (77.4% Na_2_HPO_4_, 22.6% NaH_2_PO_4_, pH 7.4), 100 mM NaCl, 0.5% NP-40, 10% glycerol, 1 mM TCEP and protease inhibitor cocktail (Roche)). After centrifugation, the supernatant was mixed with Ni-NTA agarose resin (Yeasen) at 4 °C for 2 h. The resin was washed with 20 CV lysis buffer supplemented with 15 mM imidazole and the protein was eluted with 300 mM imidazole. The eluted protein was dialyzed overnight at 4 °C in a dialysis buffer containing TEV protease to remove the tag. The protein was further purified by SEC using Superdex 200 Increase 10/300 GL (Cytiva) equilibrated with 25 mM Tris-HCl [pH 7.5], 150 mM NaCl, 1 mM TCEP and 10% glycerol.

For biochemical assays and cryo-EM sample preparation, HBT-tagged UBE3C and its mutants were expressed in Expi293 cells. 500 mL of 2×10^6^ cells ml^-1^ were transiently co-transfected with 0.5 mg UBE3C or its mutants and 0.25 mg Calmodulin plasmids using PEI (polyethylenimine). 12 h post-transfection, cells were treated with 10 mM Sodium butyrate and harvested for another 48 h. Collected cells were resuspended in a buffer (50 mM HEPES (pH 7.5), 100 mM NaCl, 0.5% NP-40, 10% glycerol, 1 mM DTT, 1mM EGTA and protease inhibitor cocktail (Roche)) and lysed by Dounce-homogenizing. The lysate was centrifuged, and the supernatant were mixed with Streptavidin affinity resin (Yeasen) at 4 °C for 2 h. The resin was wash with 50 mM Tris-HCl [pH 7.5], 100 mM NaCl, 1 mM DTT, 0.5 mM EGTA, 10% glycerol, then was incubated with TEV protease (Invitrogen) overnight at 4 °C. The flow-through was subjected to SEC on a Superdex 200 Increase 10/300 GL column (Cytiva) that was equilibrated with 25 mM Tris-HCl (pH 7.5), 150 mM NaCl, 1 mM DTT, 10% glycerol.

USP14 was recombinantly expressed in BL21-CondonPlus (DE3)-RIPL cells (Shanghai Weidi) and purified as previously reported^9^. Briefly, cells transformed with pGEX-4T-USP14 were induced by 0.2 mM IPTG and grown overnight at 20 °C. Cells were harvested and lysed in buffer containing 25 mM Tris-HCl [pH 8.0], 150 mM NaCl, 0.2% NP-40, 1 mM DTT, 10% glycerol and protease inhibitor cocktail. After clarification by centrifugation, the supernatant was incubated with pre-equilibrated Glutathione Agarose Resin (Yeasen) for 2 h at 4 °C. The resin beads were washed with buffer containing 50 mM Tris-HCl [pH 8.0], 150 mM NaCl, 10% glycerol and 1 mM DTT, followed by on-resin thrombin cleavage overnight at 4 °C in cleavage buffer (50 mM Tris-HCl [pH 8.0], 150 mM NaCl and 1 mM DTT). The eluted proteins were further purified by size-exclusion chromatography using a Superdex 75 10/300 GL column (Cytiva) pre-equilibrated with FPLC buffer (50 mM Tris-HCl [pH 8.0], 150 mM NaCl, 1 mM DTT and 10% glycerol).

### Reconstitution of UBE3C activity at the proteasome

Structural characterization of human UBE3C has historically been impeded by its propensity for aggregation following affinity chromatography. SDS-PAGE analysis of the limited monodispersed fractions recovered revealed the presence of endogenous calmodulin (Extended Data Fig. 1). We hypothesized that UBE3C aggregation is triggered by the loss of calmodulin; consistent with this, co-expression of calmodulin effectively mitigated aggregation, enabling the purification of high-concentration UBE3C. Notably, the addition of excess Ca^2+^ triggered the partial dissociation of calmodulin from UBE3C, resulting in reduced stability^44^ (Extended Data Fig. 1f, g) and renewed aggregation as observed in size-exclusion chromatography (Extended Data Fig. 1d, e). Functionally, while Ca^2+^ enhances the ubiquitin ligase activity of isolated UBE3C, it paradoxically reduces the ubiquitin-elongating activity of UBE3C when associated with the proteasome, presumably by inducing its dissociation from the proteasome complex^44^ (Extended Data Fig. 1h-k).

To prime the substrate for reconstituting UBE3C’s E4 ubiquitin-elongating activity, we performed in vitro monoubiquitylation on surface-accessible lysine residues. This was achieved using a ubiquitin mutant (with all six lysine residues mutated to alanine except for Lys48) and a WW-HECT E3 ligase derived from Rsp5 for monoubiquitylation experiments. We hypothesized that this “primed” substrate would facilitate site-specific recruitment of UBE3C to the proteasome and stabilize the E3-proteasome interaction. The substrate-conjugated monoubiquitin mutant with only one lysine residue at Lys48 exposed ensures homogeneous Lys48 linkage for the addition of second ubiquitin by UBE3C on the proteasome, although we used native ubiquitin for subsequent ubiquitin chain growth reaction.

### Liquid chromatography–tandem mass spectrometry (LC-MS/MS)

For determining the proteins bound to UBE3C, purified UBE3C from stable HEK293T cells was subjected to Liquid chromatography-tandem mass spectrometry (LC-MS/MS) assay. Proteins were reduced with 5 mM DTT and alkylated with 10 mM iodoacetamide (IAA), and then incubated with trypsin (mass ratio of 1:50) overnight at 37 °C. The enzymatic digestion was terminated by 0.1% trifluoroacetic acid (TFA). LC-MS/MS assay was performed using Easy-nLC 1200 system (Thermo Fisher), Acclaim PepMap100 C18 column (75 μm × 20 mm, Thermo Fisher) and Orbitrap Fusion Lumos mass spectrometer (Thermo Fisher). In brief, Dissolved peptide samples were loaded onto a C18 column and separated using a segmented gradient. Eluting peptides were analyzed in the data-dependent mode on a Thermo Fusion Lumos mass spectrometer. The generated data were searched against Uniprot Homo sapiens (Human) using Proteome Discoverer v.2.2 software (Thermo Fisher).

### Co-precipitation assays

For 26S proteasome co-precipitation assay, HTBH-RPN11 Expi293 cells were lysed in lysis buffer containing 50 mM HEPES (pH 7.5), 100 mM NaCl, 0.5% NP-40, 10% glycerol, 1 mM DTT and protease inhibitor cocktail (Roche), with or without the addition of 1 mM CaCl_2_ or 1 mM EGTA. The cleared cell lysates were incubated with Streptavidin affinity resin (Yeasen) at 4 °C for 2 h. Following incubation, the resin was washed with lysis buffer, and the co-precipitated proteins were eluted with 2×SDS loading buffer and subsequently analyzed by western blot using anti-RPN10 (Abcam, 1:5000 dilution), anti-UBE3C (Abcam, 1:3000 dilution) and anti-Calmodulin (Abcam, 1:3000 dilution).

For UBE3C co-precipitation assay, Expi293 cells were transiently co-transfected with UBE3C and Calmodulin plasmids at a ratio of 2:1 using PEI. After 60 h, cells were harvested and subjected to co-precipitation assay following the same protocol as described above.

### Microscale thermophoresis (MST)

The human proteasomes were labelled using the Monolith NT™ Protein Labeling Kit (NanoTemper) and exchanged to binding buffer containing 50 mM Tris-HCl [pH 7.5], 100 mM NaCl, 1 mM ATP, 5 mM MgCl_2_ and 0.05% Tween-20. To assess proteasome-UBE3C interactions, a serial 1:1 dilution series of UBE3C was prepared in the same buffer. Equal volumes of labelled proteasome were added to each UBE3C dilution, and microscale thermophoresis measurements were performed using the Monolith^TM^ NT.115 (NanoTemper) at 20% LED excitation power and 40% MST power.

### Thermal unfolding analysis

The thermal stability of UBE3C-calmodulin was evaluated using nano differential scanning fluorometry (nanoDSF). The complex was incubated for 1 h at room temperature in a buffer containing 50 mM Tris-HCl [pH 7.5], 100 mM NaCl, 1 mM DTT and varying concentrations of CaCl_2_. Intrinsic fluorescence emission at 350 and 330 nm was monitored for the complex (100 μg ml^-1^) over a temperature range of 25 to 90°C, with a ramp rate of 1°C min^-1^, using a Prometheus Panta instrument (Nano Temper). The melting temperature (T_m_) under each condition was determined by analyzing the first derivative of melting curves.

### Immunofluorescence (IF)

Hela cells plated on coverslips were washed three times with PBS and fixed with 4% paraformaldehyde for 30 min at room temperature. After three washes with PBS, cells were permeabilized with 0.1% Triton-X-100 for 10 min and blocked with goat serum in PBS for 1h at room temperature. Then cells were incubated with primary antibodies (Goat anti-RPN1 (Abcam), Rabbit anti-UBE3C (Abcam), Mouse anti-calmodulin (Abcam)) diluted 1:50 in blocking buffer overnight at 4 °C. Cells were washed with PBS three times and incubated with Alexa Fluor 555-conjugated donkey anti-goat IgG (Abcam, 1:1000 dilution) for 1h at room temperature in dark. After three washes with PBS, the cells were incubated with Alexa Fluor 488-conjugated goat anti-mouse IgG (Thermo Fisher Scientific, 1:1000 dilution) and Alexa Fluor 647-conjugated goat anti-rabbit IgG (Thermo Fisher Scientific, 1:1000 dilution) for 1 h at room temperature in dark, and then stained with 4’,6-Diamidino-2-phenylindole (Yeasen, 1ug mL^-1^). Coverslips were washed thoroughly with PBS, mounted on glass slides using Fluoromount-G (Yeasen).and examined by a confocal microscope (ZEISS LSM 900).

### Preparation of polyubiquitinated substrates

UBE1, UbcH5A and WW-HECT were recombinantly expressed in BL21-CondonPlus (DE3)-RIPL cells (Shanghai Weidi) and purified as previously reported^9^. For purification of substrates Ub4-sfGFP-NtSic1^PY^, Ubl-sfGFP-NtSic1^PY^ and NtSic1^PY^-cp8sGFP, *E. coli* BL21-CondonPlus (DE3)-RIPL cells (Shanghai Weidi) transformed with the plasmid were grown to an OD_600_ of 0.6-0.8 in LB medium and induced with 0.2 mM isopropyl β-d-1-thiogalactopyranoside (IPTG) overnight at 20 °C. Cells were collected and lysed by sonication in 50 mM Tris-HCl (pH 7.0), 150 mM NaCl, 0.2% Triton-X-100, 10% glycerol, 1 mM TCEP and protease inhibitor cocktail (Thermo Fisher Scientific). After centrifugation, the cleared lysates were incubated with pre-equilibrated Ni-NTA Resin (Yeasen) at 4 °C for 2 h. The resin was washed with 20 CV lysis buffer and 20 CV washing buffer containing 50 mM

Tris-HCl (pH 7.0), 100 mM NaCl, 20 mM imidazole, 10% glycerol and 1 mM TCEP. Proteins were then eluted with elution buffer containing 50 mM Tris-HCl (pH 7.0), 100 mM NaCl, 300 mM imidazole, 10% glycerol and 1 mM TCEP, and were further purified by SEC using Superdex 200 Increase 10/300 GL (Cytiva) equilibrated with 25 mM Tris-HCl (pH 7.0), 150 mM NaCl, 1 mM DTT, 10% glycerol.

To ubiquitylate Ub4-sfGFP-NtSic1^PY^, 1.2 μM Ub4-sfGFP-NtSic1^PY^, 0.25 μM UBE1, 2 μM UbcH5A, 1 μM WW-HECT and 20 uM K48-only Ub (Boston Biochem) were incubated in reaction buffer (50 mM Tris-HCl (pH 7.5), 100 mM NaCl, 10 mM MgCl_2_, 2 mM ATP, 1 mM TCEP, 10% glycerol) at 30 °C for 2 h. The polyubiquitinated Ub4-sfGFP-NtSic1^PY^(Ub4-sfGFP-NtSic1^PY^-Ub*_n_*) was purified by incubating with Ni-NTA resin (Yeasen) at 4 °C for 1 h. The resin was washed with 20 bed volumes of the washing buffer (50 mM Tris-HCl (pH 7.5), 300 mM NaCl, 10% glycerol, 1 mM TCEP and 25 mM imidazole). The Ub4-sfGFP-NtSic1^PY^-Ub*_n_* was eluted with the washing buffer containing 150 mM imidazole and exchanged to the storage buffer containing 50 mM Tris-HCl (pH 7.5), 100 mM NaCl, 1 mM TCEP and 10% glycerol. Since the E3 ligase WW-HECT preferentially catalyzes Lys63-linked ubiquitination^63^, The ubiquitinated substrate Ub4-sfGFP-NtSic1^PY^-Ub*_n_*is expected to only bind one monoubiquitin at each ubiquitination site. The ubiquitylation of NtSic1^PY^-cp8sGFP (Ub*_n_*-NtSic1^PY^-cp8sGFP) was performed as described above, except that wildtype ubiquitin was used.

### Ubiquitylation assay

To assess ubiquitination activity of UBE3C and its mutants, 0.25 μM UBE1, 1 μM UBE2D1 (UbcH5A), 40 µM Ub and 1 μM UBE3C or its mutants were incubated in reaction buffer (50 mM Tris-HCl (pH 7.5), 50 mM NaCl, 10 mM MgCl_2_, 5 mM ATP, 1 mM DTT) at 37 °C for 30 min. The reactions were terminated by adding 1×SDS loading buffer and subsequently subjected to western blot using anti-K29/K48/K33-linkage specific polyubiquitin antibodies (ABclonal, 1:1000-2000 dilution).

### In vitro degradation assay

Human 26S proteasomes (40 nM) were incubated with 0.4 μM UBE3C (with 0.4 μM USP14 included where indicated) for 10 min at room temperature in a degradation buffer (50 mM Tris-HCl (pH 7.5), 10 mM MgCl_2_, 50 mM NaCl, 5 mM ATP and 1mM DTT). Then 0.2 μM polyubiquitylated substrate Ub4-sfGFP-NtSic1^PY^-Ub*_n_*, 0.125 μM UBE1, 0.5 μM UBE2D1 (UbcH5A) and 20 μM Ub (in some cases K29-only Ub or K48-only Ub) were added into the mixture and incubated for 0, 2.0, 5.0, 15 and 30 min at 37 °C. The reactions were terminated by adding 1×SDS loading buffer and subsequently subjected to western blot using anti-GFP antibody (Clontech, JL-8, 1:1000 dilution).

### In cell degradation assay

UBE3C-knockout HEK293T cells stably expressing DD-sfGFP were maintained in DMEM medium supplemented with 10% FBS and 1 μM Shield (S1), a ligand that stabilizes the destabilizing domain (DD) and protects the sfGFP fusion protein from degradation^20,21,48^. Cells were transiently transfected with UBE3C variants and cultured for 24h. Following transfection, S1 was washed out to reduce degradation of DD-sfGFP, and the cells were cultured for an additional 12 h before being harvested for flow cytometry or Western blot analysis using anti-GFP and β-Tubulin antibodies.

### ATPase activity assay

ATPase activity of the human proteasome was evaluated using a malachite green phosphate assay kit (Sigma), following the manufacturer’s instructions. In brief, 30 nM proteasome and 0.3 μM UBE3C were incubated in buffer containing 50 mM Tris-HCl (pH 7.5), 100 mM NaCl, 5 mM MgCl_2_ and 0.5 mM ATP for 10 min at room temperature. 300 nM Ub*_n_*-Sic1^PY^ or

Ub*_n_*-NtSic1^PY^-cp8sGFP was added to the mixture and incubated at 37°C for 1 min. Then 20 μL Working Regent (malachite green buffer) was added to the mixture and incubated for 30 min at room temperature for color development. The absorbance at 620 nm was measured using a Varioskan Flash spectral scanning multimode reader (Thermo Fisher).

### Sample and grid preparation for cryo-EM

To assess the regulation of UBE3C on proteasomal degradation of superfolded substrates, human 26S proteasomes (0.5 μM) were incubated with 5 μM UBE3C for 10 min at room temperature. The mixture was then exchanged to an imaging buffer containing 50 mM Tris-HCl (pH 7.5), 50mM NaCl, 10 mM MgCl_2_, 5 mM ATP and 1 mM DTT. Then 1 μM UBE1, 2.5 μM UbcH5A, 150 μM Ub and 2.5 μM Ub4-sfGFP-NtSic1^PY^-Ub*_n_*were added to initiate the degradation reaction, which was incubated for 1, 2, 5 and 15 min at 37 °C. For each time point, the reaction mixture was treated with 0.005% NP-40 and applied to glow-discharged grids (Quantifoil, R1.2/1.3, Au, 300 mesh) for immediate cryo-plunging. The grids were blotted for 1.5 s at 4 °C with 100% humidity and plunged into liquid ethane cooled by liquid nitrogen using a Vitrobot Mark IV (Thermo Fisher Scientific). To assess the allosteric regulation of proteasome-bound UBE3C and USP14, 0.5 μM proteasomes were incubated with 5 μM UBE3C and 5 μM USP14 for 10 min at room temperature. The mixture was then exchanged to an imaging buffer containing 50 mM Tris-HCl [pH 7.5], 100 mM NaCl, 10 mM MgCl_2_, 5 mM ATP and 0.5 mM DTT. Then 1 μM UBE1, 2.5 μM UbcH5A, 20 μM Ub and 5 μM

Ub*_n_*-NtSic1^PY^-cp8sGFP were added to initiate the degradation reaction, which was incubated for 15, 30, 60 and 120 s at 37 °C. For each time point, 0.005% NP-40 was added to the reaction mixture before following the same cryo-plunging protocol as described above.

### Cryo-EM data collection

Cryo-grids were initially screened using a 200 kV Talos Arctica microscope (Thermo Fisher). High-quality grids were then transferred to a 300 kV Titan Krios G2 microscope (Thermo Fisher), equipped with a post-column BioQuantum energy filter (Gatan) connected to a K3 direct electron detector (Gatan). Coma-free alignment and parallel illumination were manually optimized before each data collection session. Cryo-EM data were automatically acquired using SerialEM software^64^ in super-resolution counting mode with 20 eV energy slit. The nominal defocus was set within the range of-0.8 to-2.0 µm. The data collection was performed with a super-resolution pixel size of 0.685 Å and an accumulated dose of ∼50 electrons per Å². For the UBE3C-proteasome dataset, 15,215, 15,899, 13,297, 8,647 movies were collected for cryo-grids made with the reaction time of 1min, 2min, 5min, 15 min, respectively. For the USP14-UBE3C-proteasome dataset, 7,922, 14,375, 8,218, 7,808, 3,617 movies were collected for cryo-grids made with the reaction time of 15s, 30s, 1min, 2min, 5min, respectively.

### Cryo-EM data processing

Drift correction was performed using RELION^65^’s own implementation of the MotionCor2 program^66^, with a super-resolution pixel size of 0.685 Å. The CTF parameters of the drift-corrected micrographs were all calculated using the Gctf program^67^. Particles were automatically picked using an improved version of the DeepEM program^68^ on micrographs that were fourfold binned to a pixel size of 2.74 Å. Micrographs screening and auto-picked particles checking were then performed using EMAN2 software^69^. For the 1min, 2min, 5min and 15 min subsets of the UBE3C-proteasome dataset, a total of 1,629,150, 1,727,709, 1,382,191, and 776,299 particles were picked, respectively. For the 15s, 30s, 1min, 2min, 5min subsets of the USP14-UBE3C-proteasome dataset, a total of 706,015, 2,154,438, 952,224, 1,009,318, 396,507 particles were picked, respectively. Reference-free 2D classification and 3D classification were carried out using RELION^65^ version 4.0 and ROME^70^. Focused 3D classification, CTF and aberration refinement, as well as high-resolution refinement, were performed with RELION^65^, and the AlphaCryo4D software^71^ was used to analyze conformational changes and conduct in-depth 3D classification.

Both datasets employed a similar hierarchical 3D classification strategy for data analysis (Extended Data Figs. 2, 12), optimized as described previously^10^. The main data analysis process for both datasets was very similar. The data analysis process for UBE3C-proteasome dataset will be described in detail, while for USP14-UBE3C-proteasome dataset, the focus will primarily be on the differences from the first dataset, for the sake of brevity.

For the UBE3C-proteasome dataset, subsets from different conditions were processed separately at steps 1 and 2, and combined at steps 3, 4 and 5. In the focused 3D classification, the same dataset was classified separately based on the conformation of proteasome and UBE3C. Then the intersection of the two classification results was taken to obtain combinations of different proteasome and UBE3C states.

*Step 1*: Doubly capped proteasome particles were separated from singly capped particles through several rounds of 2D and 3D classification. These particles were then aligned to the consensus models of the doubly and singly capped proteasome to obtain their approximate shift and angular parameters. Using these parameters, each doubly capped particle was split into two pseudo-singly capped particles by re-centering the box onto the RP–CP subcomplex. The box size of both the pseudo-singly capped and true singly capped particles was then reduced to 640 × 640 pixels with a pixel size of 0.685 Å, followed by a two-fold down-sampling to a pixel size of 1.37 Å for subsequent processing. After this step, a total of 5,980,852 particles from all subsets were obtained.

*Step 2*: Particles were aligned to the CP subcomplex through auto-refinement, followed by an alignment-skipped RP-masked 3D classification, which separated the E_A_-like states from the E_D_/S_D_-like states. Poor 3D classes containing partially broken RP were removed for further analysis at this step. A total of 2,079,616 E_A_-like particles and 2,835,215 E_D_/S_D_-like particles were obtained.

*Step 3*: To improve the accuracy and quality of the 3D classification, we combined the E_A_-like states and E_D_/S_D_-like states from subsets of different conditions into separate clusters for further processing. For the two clusters, CP-masked auto-refinement was performed, followed by CTF and aberration refinement. For the E_D_/S_D_-like states, another run of CP-masked auto-refinement was performed, after which the CP density was subtracted, and then the particle box was recentered onto the RP subcomplex and shrunk to 240 × 240 pixels, with a pixel size of 1.37 Å. For the E_A_-like states, since the binding rate of UBE3C was quite low and analysis of E_A_-like states was relatively straightforward, subtraction was omitted to save time. Both clusters then underwent multiple rounds of RP-masked auto-refinement and alignment-skipped RP-masked 3D classification. Due to the complexity of the E_D_/S_D_-like states, AlphaCryo4D analysis^71^ was also used to assist with their conformational analysis and classification. Ultimately, 14 proteasome states were obtained, named EA1^UBE3C^, EA2.1^UBE3C^, EA2.2^UBE3C^, ED0^UBE3C^, ED1.1^UBE3C^, ED1.2^UBE3C^, E_D1.8_^UBE3C^, E_D1.9_^UBE3C^, E_D2.1_^UBE3C^, E_D2.2_^UBE3C^, E_D3.1_^UBE3C^, E_D3.9_^UBE3C^ S_B_UBE3C, further classification was performed to improve the density quality in regions such as pore loops.

*Step 4*: For the E_D_/S_D_-like states, CP-subtracted particles from the previous step were reused for UBE3C conformational classification. All particles from the E_D_/S_D_-like states were also combined here and subjected to subsequent refinement and classification in this step. After

RP-masked auto-refinement, although the intact UBE3C density was weak, the density of its N-terminus was almost as strong as that of the nearby proteasome subunits. After multiple classification attempts, we were unable to separate the particles with the UBE3C N-terminus bound from those without it, suggesting that most of the E_D_/S_D_-like particles have the UBE3C N-terminus bound. Several rounds of UBE3C-masked classification were performed, after which particles with intact UBE3C and those with only the N-terminus bound were separated. A total of 1,234,576 particles with intact UBE3C binding were obtained, and then they were aligned to the Lid and UBE3C, followed by multiple rounds of UBE3C-masked skip-alignment 3D classification and AlphaCryo4D analysis^71^. After excluding low-resolution classes where secondary structure was unclear, we obtained four high-resolution states, named P1, P2, P3, and P4. Only a single ubiquitin density was well-resolved in states P1 and P2. In contrast, distinct densities for three and four ubiquitin moieties were clearly observed in P3 and P4, respectively. However, upon low-pass filtering and visualization at a lower contour level, five ubiquitin moieties became apparent in both the P3 and P4 maps.

For the E_A_-like states, particles were combined and classified based on the UBE3C N-terminus, revealing that—unlike the E_D_/S_D_-like states—less than 12% of particles exhibited a bound N-terminus. The particles with the N-terminus bound then underwent UBE3C-masked skip-alignment 3D classification, resulting in 137,434 particles with intact UBE3C binding.

Further classification excluded classes with poorly resolved secondary structures, retaining four relatively good classes. Following UBE3C-masked refinement, the local resolution of UBE3C was limited to around 6 Å for all four classes. Two of these classes broadly resembled the UBE3C P1 and P3 states identified in the E_D_/S_D_-like states and were labeled accordingly. The remaining two classes exhibited significant heterogeneity and could not be unambiguously assigned to the UBE3C states observed in the E_D_/S_D_-like states; these were therefore termed Mix1 and Mix2. Due to the limited particle number and low resolution, the four UBE3C classes of E_A_-like states were not subjected to final refinement.

*Step 5*: Time-resolved analysis of conformational changes was performed for the E_A_-like states, E_D_/S_D_-like states, and UBE3C states by simply separating particles based on their time labels. The proportion of particles of each state at a given time point was calculated by summing the number of particles for all states at the same time point and then calculating the fraction of particles of each state with respect to the total number of particles at this time point^72,73^.

Subsequently, for the particles belonging to the 11 E_D_/S_D_-like states and the four UBE3C states, intersections were calculated to determine the combinations of different proteasome and UBE3C states. For the E_D_/S_D_-like states, particles used for this analysis were taken from the classification results prior to the further classification aimed at improving pore loop density.

All sub-datasets corresponding to the 11 E_D_/S_D_-like proteasome states are observed to each comprise all four UBE3C states. In other words, any of the 11 proteasome states is observed to coexist with any of the four UBE3C states in a subset of the cryo-EM dataset. Thus, there are in total 44 (i.e., 4 times 11) approximately homogenous conformers of UBE3C-proteasome complex identified in the entire dataset that are contributed to the final high-resolution reconstructions, although such 3D classification does not exclude the potential existence of other lowly populated conformational states that are insufficient for high-resolution refinement, which we intent to exclude in the following structural analysis and discussion. Among the resulting 44 conformers, four conformers with both a high particle number and good map quality—ED1.8^UBE3C^ ^P1^, ED2.1^UBE3C^ ^P1^, ED1.8^UBE3C^ ^P4^, ED2.1^UBE3C^ ^P4^—were selected for final refinement.

Final refinements for all states were performed using two-fold binned particles with a pixel size of 1.37 Å. Different focused refinement strategies were applied to the 14 individual proteasome states, the four individual UBE3C states of the E_D_/S_D_-like dataset, and the four selected intersected conformers. For individual proteasome states and the intersected conformers, refinement was conducted on pseudo-singly capped particles. For individual proteasome states, RP-masked and CP-masked refinements were performed for each state, and based on the results of auto-refinement, half-maps were reconstructed using the original particles with a pixel size of 0.685 Å. For individual UBE3C states, UBE3C-masked, HECT domain-masked, and Lid subcomplex-masked refinements were performed separately for each state. For intersected conformers ED1.8^UBE3C^ ^P1^ and ED2.1^UBE3C^ ^P1^, focused refinements were carried out using CP, RP, UBE3C, and HECT masks. For conformers ED1.8^UBE3C^ ^P4^and ED2.1^UBE3C^ ^P4^, the HECT mask was replaced with a C-lobe mask, as tests showed that using the C-lobe mask yielded higher local resolution for the P4 combinations, especially when they contained fewer particles. Refinement of the intersected conformers indicated that, in the resulting maps, the proteasome and UBE3C regions were structurally consistent with their corresponding parental states. To facilitate the visualization of structural details, a composite map generated from the higher-resolution maps of the parental proteasome and UBE3C states is presented in Fig. 1b to represent the intersected conformer. Fourier shell correlation (FSC) curves for all 14 proteasome states were measured in a gold-standard procedure, yielding nominal resolutions ranging from 2.5 to 3.6 Å and local RP resolutions ranging from 2.7 to 4.1 Å. The FSC curves for all four UBE3C states were also measured using the same procedure, with local UBE3C resolutions ranging from 3.8 to 4.4 Å. Before visualization and final model refinement, B-factor sharpening was applied to each set of focused refinement results, followed by map combination using Phenix^74^ to generate composite maps for each state.

To further improve the local density quality of the ubiquitins in state P4, additional classification was performed using a mask covering the C-lobe and ubiquitins. 3D classes with blurred Ub^D^ and Ub^A^ densities were excluded, and class with improved ubiquitin densities including 57,661 particles were then selected and refined. The locally improved maps were used only for visualization and for adjusting the atomic models of the C-lobe and ubiquitins.

For the USP14-UBE3C-proteasome dataset, the processing workflow is highly similar to that used for the UBE3C-proteasome dataset. Steps 1 and 2 were virtually identical to the previous pipeline. Step 1 yielded 5,830,247 particles, comprising both pseudo-and truly single-capped proteasomes. Following Step 2, the dataset was separated into 2,989,171 E_A_-like particles and 1,952,336 E_D_/S_D_-like particles. In Step 3, signal subtraction was also applied to E_A_-like particles to improve classification precision and probe for potential USP14-bound conformations. However, the focused classification did not reveal distinct USP-bound conformations. It resolved three states, named EA1, EA2.2 and EA2.2^UBE3C^, with states EA2.2 and E_A2.2_^UBE3C^ exhibiting high structural similarity throughout the entire RP. For the E_D_/S_D_-like particles, classification of the RP yielded conformations E_Do_^UBE3C^, E_D1.1_^UBE3C-UBL^, ^E^D2.1^UBE3C-USP14^, ^E^D3.1UBE3C-USP14, ^E^D3.9^UBE3C-USP14^, S_B_^UBE3C^, ^S^D4^UBE3C-USP14^. Apart from the USP14, these structures were virtually indistinguishable from those observed in the UBE3C-proteasome dataset and are therefore not described individually (correspondence shown in Extended Data Fig. 12a).

A key difference in Step 3 was that, following RP classification, each E_D_/S_D_-like state underwent focused 3D classification targeting the USP domain. However, for the majority of the RP states, characterizing the USP domain proved challenging. Restricted by limited particle numbers and intrinsic conformational flexibility, the USP domain densities were generally ill-defined, resulting in either unsuccessful classification or insufficient local resolution.

Specifically, no clear USP14 density was observed for states Eo^UBE3C^ and S_B_^UBE3C^, while only the UBL domain density was visible in E_D1._^UBE3C-UBL^. For states such as E_D3.1_^UBE3C-USP14^,although USP binding was evident in certain classes, the USP domain could not be resolved to high resolution. Ultimately, only the state with the largest particle population, E_D2.1_^UBE3C-USP14^ successfully yielded three high-resolution USP14-bound conformations, named E_D2.1_^UBE3C-UBL^, E_D2.1_^UBE3C-USP14^ and E_D2.1_^UBE3C-USP14-Ub^. These three states were subjected to final refinement using the same protocol as the proteasome states of the UBE3C-proteasome dataset, achieving nominal resolutions of 2.8 Å, 3.2 Å, and 3.1 Å, and RP-masked resolutions of 3.1 Å, 3.5 Å, and 3.4 Å, respectively.

In Step 4, given the low binding rate of UBE3C on E_A_-like particles, no further classification of UBE3C conformations was performed. For all E_D_/S_D_-like particles, initial classification identified 525,546 particles containing intact UBE3C. Following multiple rounds of focused 3D classification, two states, P1^USP14^ and P3^USP14^, were resolved. As these states were structurally consistent with the P1 and P3 states from the UBE3C-proteasome dataset, they are not described further.

### Atomic model building and refinement

To build the atomic models of the individual UBE3C states, the AlphaFold 3^75^-predicted structures of UBE3C and calmodulin, four copies of a published model of ubiquitin, along with the model of the lid subcomplex extracted from the published state E_D2.1_ structure^9^(PDB:7W3H) were first docked into the map of state P4 using Chimera^76^. These models were then manually rigid-body fitted block by block in Coot^77^. Published crystal structures of the HECT domain and calmodulin were also used as references. After an initial round of real-space refinement in Phenix^74^, the P4 model was rigid-body fitted into the maps of the other three UBE3C states. For each state, the corresponding fitting of ubiquitin was adjusted based on the map density, followed by manual improvements of the whole model in Coot^77^ and rounds of real-space refinement in Phenix^74^.

For the proteasome state models, published proteasome coordinates were used as the initial references. For each state, the model of the closest conformation was first docked into the map, followed by chain-by-chain manual rigid-body fitting in Coot^77^. Subsequently, for chains with minor conformational changes, such as the RPN subunits, rigid-body fitting was performed domain by domain. For chains with significant conformational rearrangements, such as the RPT subunits, rigid-body fitting was carried out in smaller segments. For all E_D_/S_D_-like states and E_A2.2_^UBE3C^, residues 1–50 of UBE3C were added based on the refined UBE3C model and the map density. Since the substrate in the central channel is not sequence-specific, polypeptide chains without assigned amino acid sequences were used to model the substrate density. The nucleotide-binding states of the ATPases were determined by unbiased fitting of ATP or ADP molecules based on the map densities in the nucleotide-binding pockets. A state was classified as ‘apo-like’ if the densities were weak and insufficient to accommodate a nucleotide molecule.

Subsequent model refinement, including real-space refinement in Phenix^74^ and manual adjustments in Coot^77^, was iterated until the model quality met the expected level.

For the three USP14-bound E_D2.1_ states, the atomic model of E^UBE3C^ from the UBE3C-proteasome dataset served as the initial template. The coordinates for USP14 and ubiquitin were derived from the previously published E_D2.1_ structure^9^ (PDB ID: 7W3H). These components were segmented into independent rigid bodies and manually docked into the density maps using Coot^77^, with regions lacking supporting density truncated specifically for each state. The models were then subjected to iterative cycles of real-space refinement in Phenix^74^ and manual adjustment in Coot^74^ to optimize model geometry. Modeling of the substrate and nucleotides was performed following the same protocol as described for the proteasome states.

### Structural analysis and visualization

All structures were analyzed using Coot^77^, PyMOL^78^, UCSF Chimera^76^, and ChimeraX^79^. Inter-subunit interactions and interfacial areas were computed and analysed using the PISA server^80^ (https://www.ebi.ac.uk/pdbe/prot_int/pistart.html). Local resolution variations were estimated with ResMap^81^. Structural figures were generated in PyMOL^78^, ChimeraX^79^, or Coot^77^. Structural alignment and comparisons were performed in PyMOL^78^ and ChimeraX^79^.

Comparisons to protein structures from previous publications used the atomic models in the PDB under accession codes: 4LZC (Apo calmodulin-bound IQCG^82^), 6MC9 (Ca^2+^-bound calmodulin with Nav1.4 C-Terminal^83^), 3OLM (ubiquitin-bound HECT domain of RSP5^84^), 2XBB (ubiquitin-bound HECT domain of NEDD4^52^), 1C4Z (E6AP-UbcH7 complex^85^), 3JVZ (NEDD4L-HECT-UbcH5B-ubiquitin complex^49^), 8C07 (ubiquitin-bound HECT domain of UBR5^50^), 7W37(state EA1^UBL^ of USP14-bound human proteasome^9^), 7W38 (state EA2.0^UBL^), 7W39 (state EA2.1^UBL^), 7W3A (state ED4^USP^^14^), 7W3C (state ED0^USP^^14^), 7W3F (state ED1^USP^^14^), 7W3G (stateE_D2.0_^USP^^14^), 7W3H (state E_D2.1_^USP14^), 7W3I (state S_B_^USP14^), 7W3K (state S_B_^USP14^).

## Author Contributions

S.Z., D.Y. and Y.M. conceived this study. S.Z designed and purified the substrates for degradation assays. S.Z. and L.Z. purified the other proteins and conducted the biochemical experiments. M.B. and D.F. contributed to the protein purification approach. D.Y. and S.Z. prepared cryo-EM samples, collected cryo-EM data and analyzed the experimental cryo-EM datasets. D.Y. refined the density maps, built the atomic models and analyzed the data. Y.M., S.Z. and D.Y. wrote the manuscript. Y.M. supervised the project.

## Acknowledgements

We thank Y. Saeki for the plasmids expressing Sic1^PY^ and WW-HECT; L. Huang for the proteasome-expressing cell line; J. Sodroski and C. Li for the Expi293 cell line and Hela cell line; C. Tang for the plasmid expressing UBE1; X. Li, Z. Guo, X. Pei, C. Fan and Y. Ma for technical assistance with cryo-EM data collection and storage; D. Liu and Q. Zhang for technical assistance with the mass spectrometry experiments. The cryo-EM data were collected at the Cryo-EM Core Facility Platform and Laboratory of Electron Microscopy at Peking University. The data processing was supported by the High-Performance Computing Platform of Peking University.

The mass spectrometry data were collected at the National Center for Protein Sciences at Peking University. This work was supported in part by National Natural Science Foundation of China (12125401 and 12090051 to Y.M. and 32471308 to S.Zou), National Key Research and Development Program of China (2023YFF1204400 and 2023YFF1204401 to Y.M.), Beijing Natural Science Foundation grant (Z180016/Z18J008 to Y.M.) and by AI for Science (AI4S)-Preferred Program at Peking University Shenzhen Graduate School.

## Data availability

LC–MS/MS raw data for purified human UBE3C have been deposited in PRIDE with the accession code PXD071142. Atomic coordinates and cryo-EM structures are being deposited in the Protein Data Bank (and the EMDB) with accession codes released upon the publication of this study. All raw data are available upon reasonable request.

---

**Extended Data Fig. 1.**
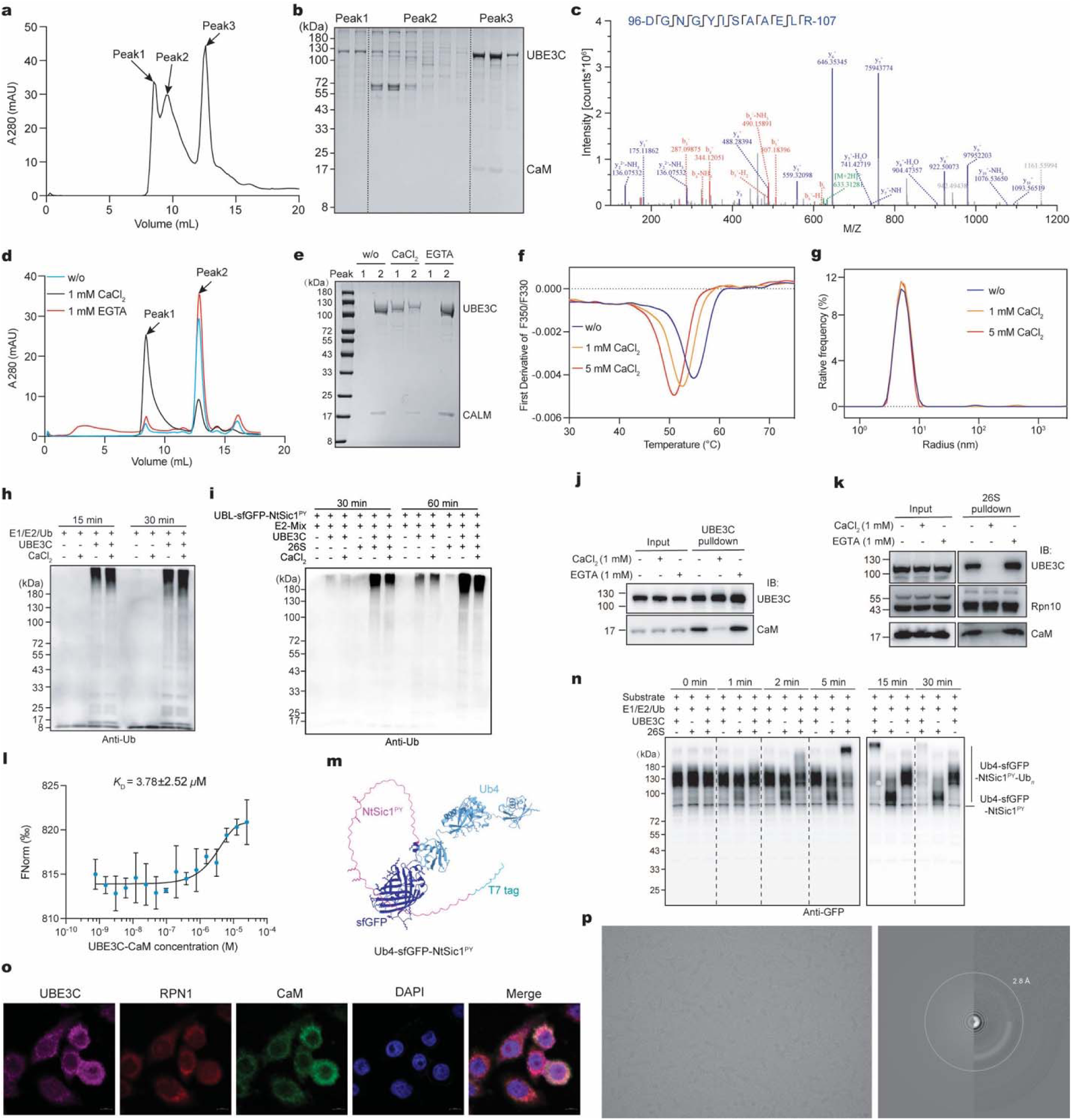
Purification and functional characterization of UBE3C, and cryo-EM imaging of USP14-proteasome complex. **a**, Purification of human UBE3C from stable HEK293T cells by size-exclusion chromatography (SEC; Superdex 200 Increase 10/300 GL). **b**, SDS-PAGE analysis of SEC fractions from **a**. **c**, LC-MS/MS analysis of purified UBE3C. **d**, SEC analysis of the UBE3C-calmodulin complex in the presence of EGTA or CaCl_2_ (repeated twice). **e**, SDS-PAGE of SEC fractions corresponding to **d**. **f**, Representative thermal denaturation curves of UBE3C-calmodulin complex in the presence of CaCl_2_. The minima of the derivative curves indicate melting temperatures (Tm) (repeated three times). g, Dynamic light scattering (DLS) analysis of UBE3C-calmodulin complex. The hydrodynamic diameter of UBE3C-calmodulin complex remained unchanged across different concentrations of CaCl_2_ (repeated three times). **h, i**, *In vitro* ubiquitylation activity of UBE3C in the presence of CaCl_2_ **(h)** or CaCl_2_ plus proteasome **(i)**. Ubl-sfGFP-NtSic1^PY^, was used to induce transient proteasome states and enhance UBE3C recruitment. Reactions were analyzed by SDS-PAGE and immunoblotting with an anti-ubiquitin antibody. A representative result from three independent experiments is shown. **j, k**, Co-precipitation assays assessing the effect of EGTA or CaCl_2_ on the interaction between UBE3C and calmodulin **(j)**, and between UBE3C and proteasome **(k)**. Co-precipitated proteins were analyzed by Western blot. Representative blot from three independent experiments. **l**, Microscale thermophoresis (MST) analysis of UBE3C-calmodulin binding to human proteasome. Dissociation constant was calculated from three independent experiments (shown as mean ± s.d.). **m**, Predicted model of the substrate Ub4-sfGFP-NtSic1^PY^ genetated using Alphafold 3^75^. **n**, In vitro degradation of a superfolder GFP-based substrate by UBE3C-human proteasome complex in the presence of E1, E2 and Ubiquitin, analyzed by western blot using an anti-GFP antibody. Substrate, Ub4-sfGFP-NtSic1^PY^-K48Ub*_n_*. Data are representative of three independent experiments. **o**, Immunofluorescence imaging for assessing UBE3C, calmodulin and proteasome. Nucleus were stained with DAPI. Scale bar, 10 μm. Images represent two independent experiments. **p**, Representative motion-corrected cryo-EM micrographs of the UBE3C-proteasome complex (left) from the time-resolved dataset at a reaction time point of 1 min. Power spectrum evaluation of the corresponding micrographs analyzed using GCTF^67^ (right).

**Extended Data Fig. 2.**
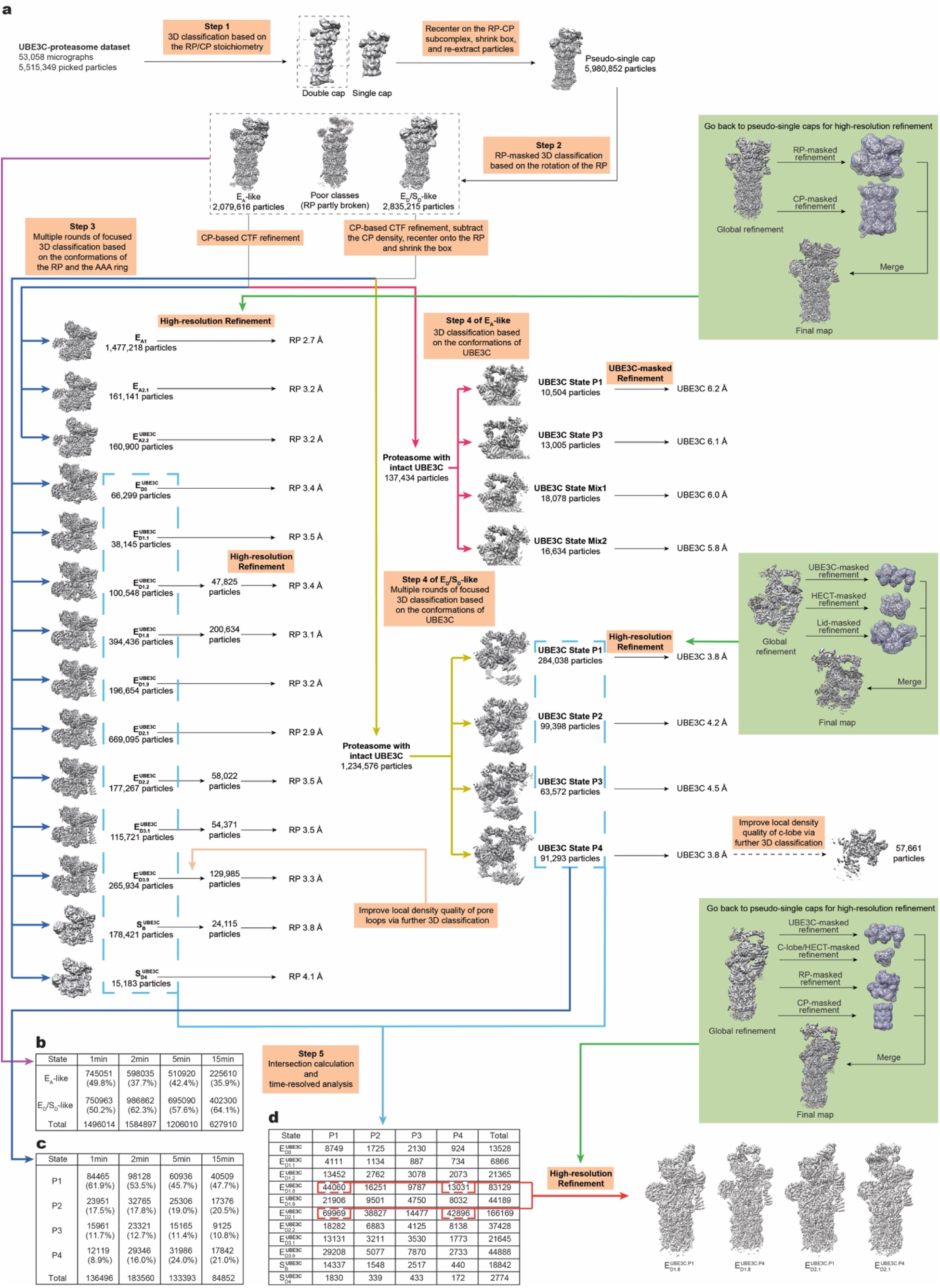
Cryo-EM data processing for the UBE3C-proteasome complex. **a**, The workflow diagram illustrates the key steps in our hierarchical 3D classification strategy. For clarity, the detailed iterations of classification at each step are omitted. The particle number following 3D classification and the final reconstruction, as well as the resolutions of the RP-masked and UBE3C-masked reconstructions for each state, are indicated. Three green boxes highlight the different focused refinement strategies used for the three kinds of states: individual proteasome states, individual UBE3C states, and the intersected conformers. **b-c**, Time-resolved analysis of proteasome and UBE3C states by restoring the time labels. Percentages were calculated based on the total particle count corresponding to a given time point. **d**, Intersection calculation results of the 9 E_D_-like states and 4 UBE3C states. Four intersected conformers with relatively large particle numbers and good map quality were selected for final reconstruction.

**Extended Data Fig. 3.**
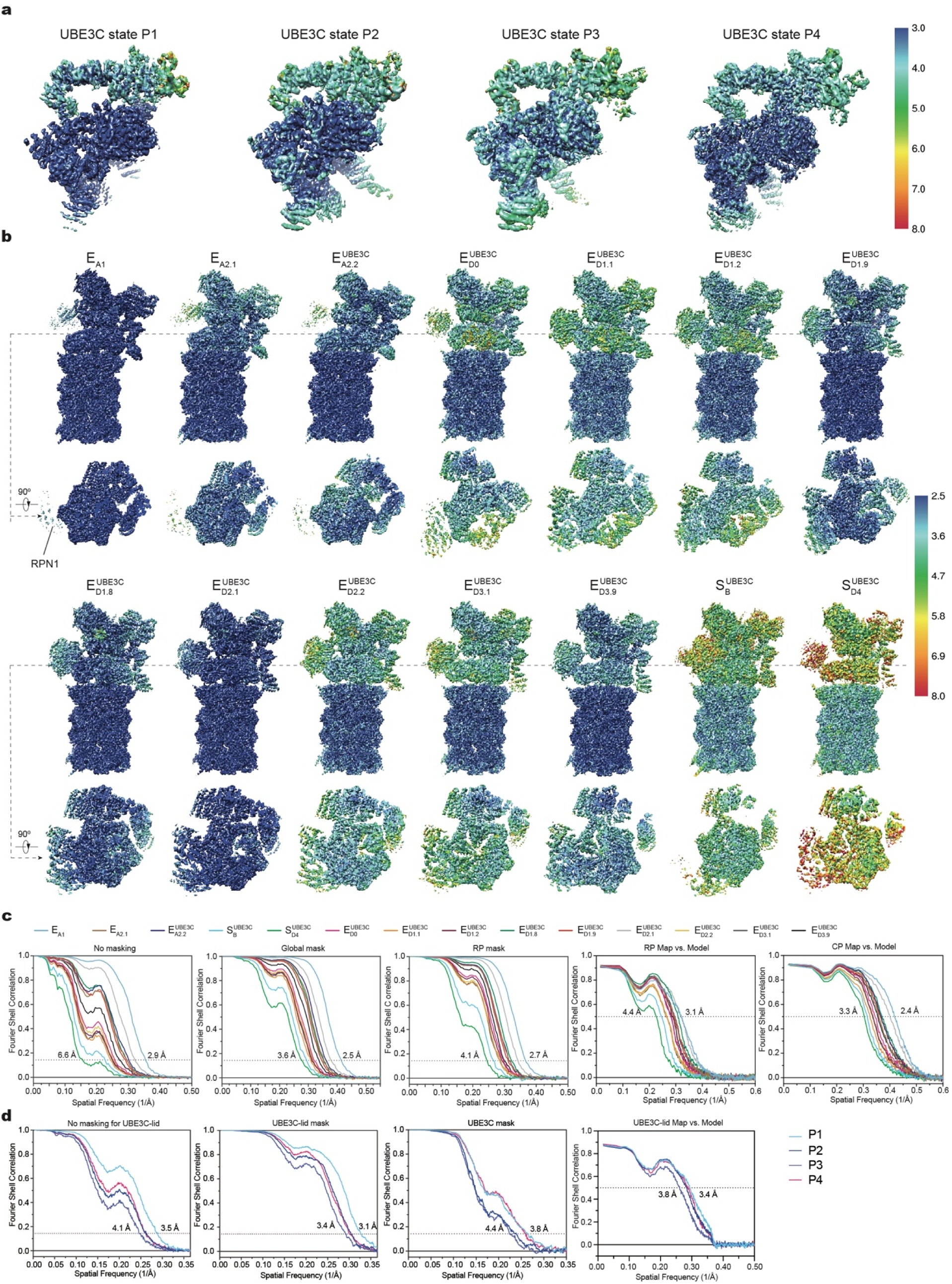
Cryo-EM reconstructions and resolution measurements of the UBE3C-proteasome complex. a, Local resolution estimation of the Lid-UBE3C reconstructions for the four UBE3C states. **b,** Local resolution estimation of the reconstructions for 14 proteasome states. Each state is presented in two orthogonal views: a side view and a cross-sectional view at the AAA domains. **c-d,** Gold-standard Fourier shell correlation (FSC) plots of the complete RP-CP maps of all proteasome states calculated without **(c)** or with **(d)** masking the separately refined half-maps. **e,** Gold-standard FSC plots of the RP-masked reconstructions of all proteasome states. The RP maps were refined by focusing the mask on the RP subcomplex. **f-g,** Model-map FSC plots calculated by Phenix between each proteasome model and its corresponding RP-masked **(f)** or CP-masked **(g)** reconstructions. **h-i,** Gold-standard FSC plots of the Lid-UBE3C maps of all UBE3C states calculated without **(h)** or with **(i)** masking the separately refined half-maps. **j,** Gold-standard FSC plots of the UBE3C-masked reconstructions of all UBE3C states. The UBE3C maps were refined by focusing the mask on the UBE3C. **k,** Model-map FSC plots calculated between each Lid-UBE3C map and its corresponding model. For each state, seperately refined Lid, UBE3C, HECT domain maps were merged in Fourier space into a single Lid-UBE3C map, which was used for the model-map FSC calculation.

**Extended Data Fig. 4.**
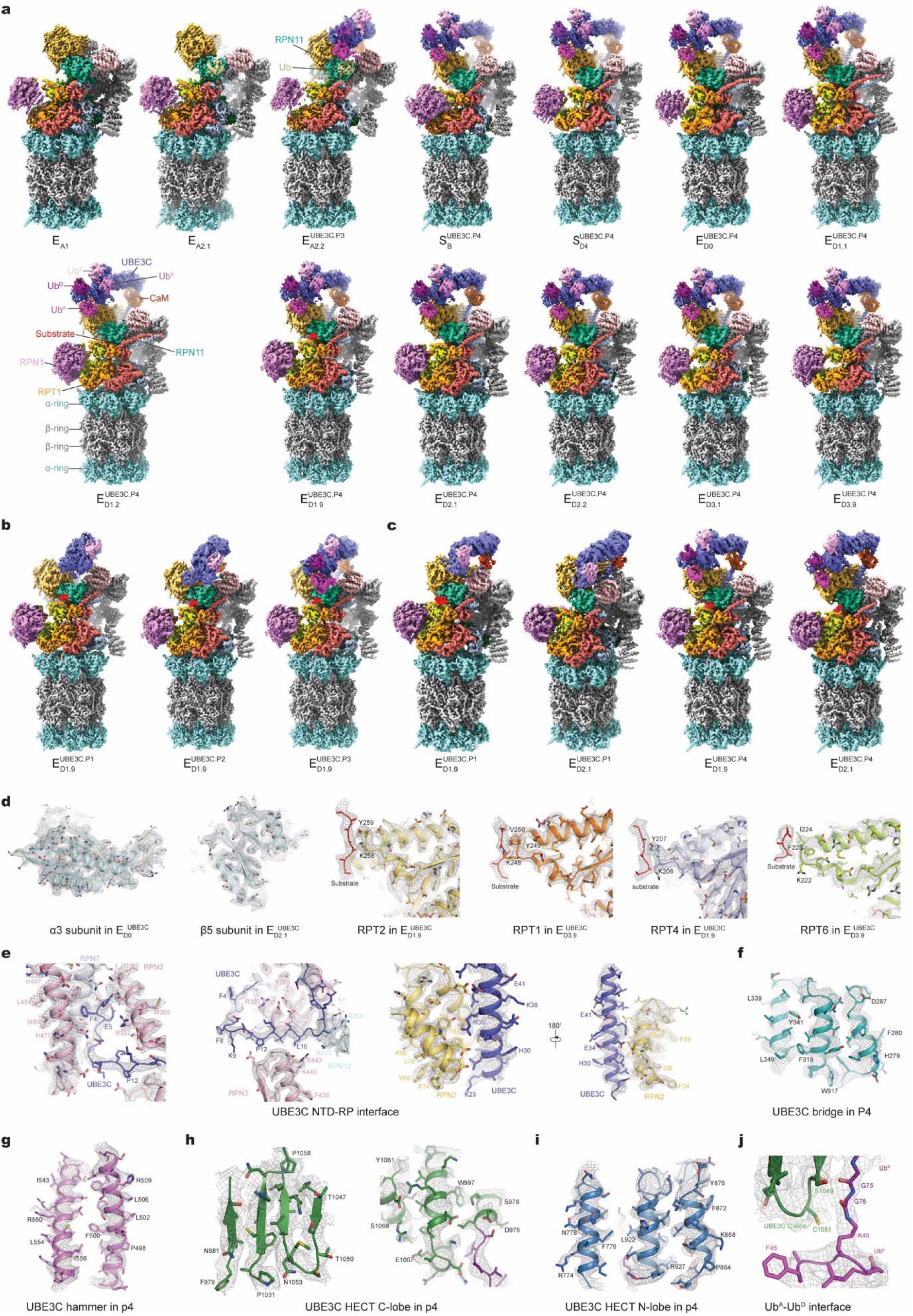
Cryo-EM maps of the UBE3C-proteasome complex. a, Gallery of refined cryo-EM maps of the proteasome. All nine E_D_-like maps, S_D_-like maps and E^UBE3C^are shown as composite reconstructions, with CP and RP densities from focused refinements and UBE3C density from the P4 state, except for E^UBE3C^, which uses the P3 state. **b,** Composite maps of the E^UBE3C^ proteasome state combined with UBE3C states P1, P2 and P3. **c,** Refined cryo-EM maps of the four intersected proteasome-UBE3C conformers identified through intersection analysis of the nine E_D_-like proteasome states and the four UBE3C states. **d-j,** Typical high-resolution cryo-EM densities (mesh) of secondary structures superimposed with their atomic models. Different subunits of the proteasome are shown in **(d)**, where the substrates shown in the right four panels are modelled using polypeptide chains without assignment of amino acid sequence. **e,** High-resolution cryo-EM densities of the UBE3C NTS binding pocket in UBE3C state P4. **f,** High-resolution cryo-EM densities of the UBE3C bridge domain in UBE3C state P4. **g,** High-resolution cryo-EM densities of the UBE3C hammer domain in UBE3C state P4. **h-i,** High-resolution cryo-EM densities of the C-lobe **(h)** and N-lobe **(i)** of the UBE3C HECT domain in UBE3C state P4. **j,** High-resolution cryo-EM densities of the Ub^D^-Ub^A^ interface in UBE3C state P4.

**Extended Data Fig. 5.**
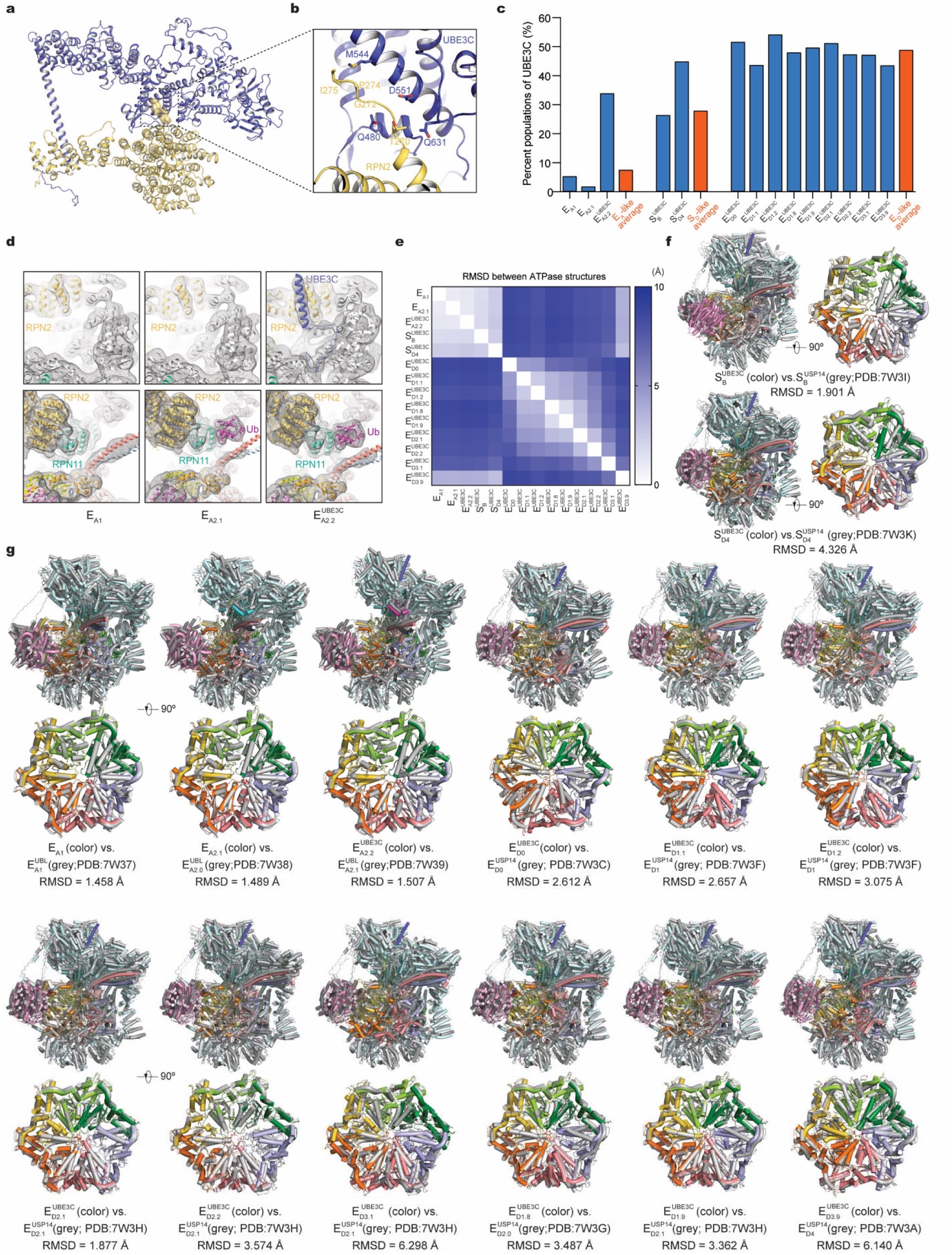
Structural comparison of UBE3C-proteasome complex with previously reported structures. a, Atomic model of the UBE3C hammer domain bound to the C-terminal domain of RPN2 in state P4. The loop above the RPN2 T1 site (residues 269–275) is highlighted in both cartoon and surface representations. **b**, Magnified view of the UBE3C–RPN2 interface shown in **a**. **c,** Quantification of UBE3C CTD binding across S_A_-like, E_D_-like, and S_D_-like states. The presence of CTD density reflects binding of full-length UBE3C. Denominators represent the total particle numbers of all sub-states within each state category. **d,** Structural comparison of states E_A1_, E_A2.1_ and E^UBE3C^ shows that the UBE3C NTD engages the proteasome specifically in the E^UBE3C^ state. Cryo-EM densities are displayed as grey mesh and low-pass-filtered to 8 Å. **e,** Root-mean-squared-deviation (RMSD) values mapped onto the AAA-ATPase ring illustrate structural differences among the proteasome states. **f, g,** Comparison of the RP and ATPase structures among S_D_-like states **(f)** and E_A_-like/E_D_-like states **(g)** with previously reported cryo-EM structures. The RMSD values for the ATPase are shown below each panel.

**Extended Data Fig. 6.**
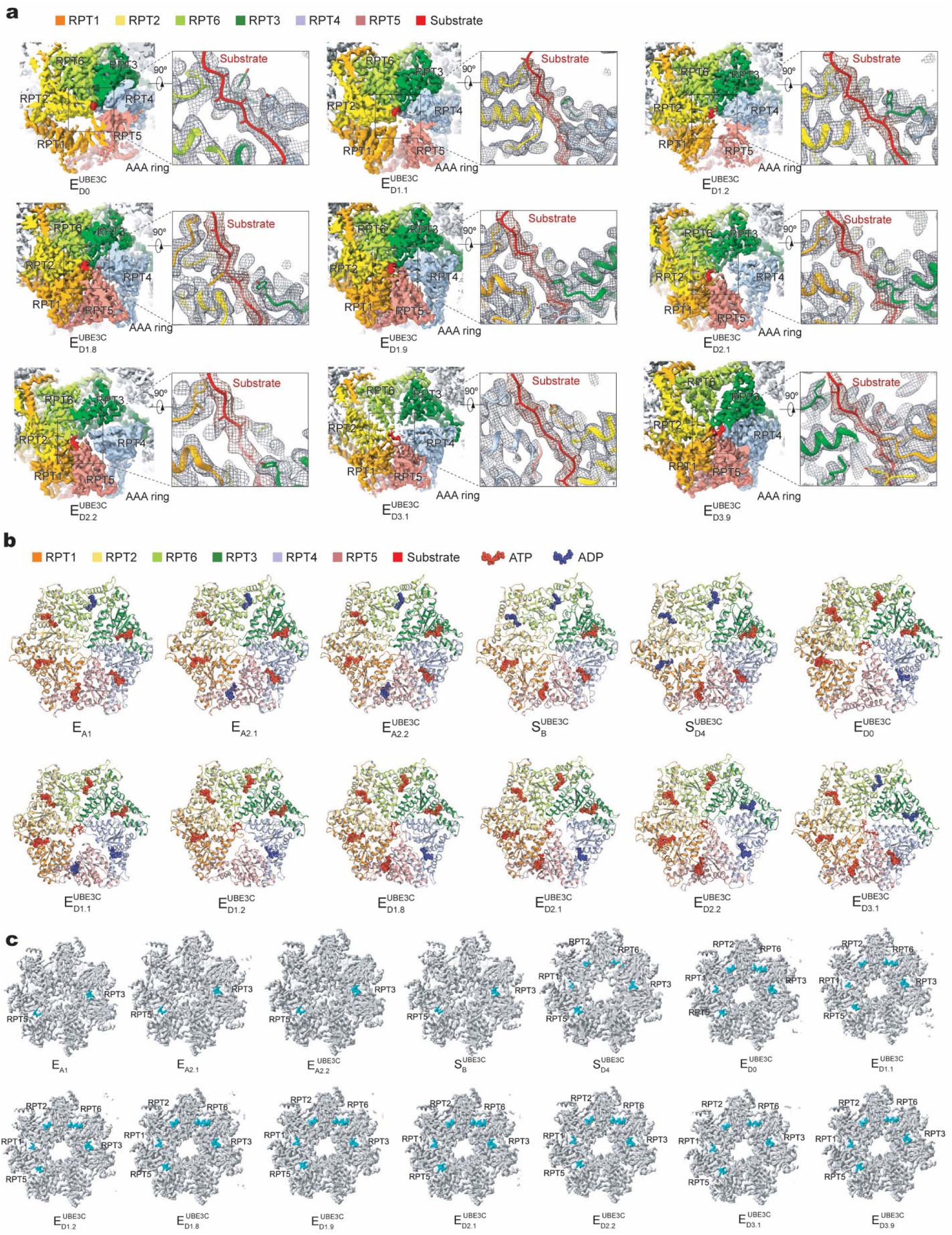
Key structural features of UBE3C-proteasome complex across different states. **a**, Cryo-EM densities of the AAA-ATPase ring bound to substrate in ED-like states. Top views of the AAA-ATPase ring (left insets) and zoomed-in side views of substrate engagement (right insets) are rendered as surface and mesh, respectively. Substrate densities are modelled as polypeptide backbones and colored in red. **b**, Top views of the AAA-ATPase ring atomic models from all resolved states, displayed as cartoon representations. Bound nucleotides are shown as spheres: ADP in blue and ATP in red. **c**, Cryo-EM densities of the RP-CP interface for all states, shown in top view. The C-terminal tails of RPT subunits inserted into the α-pockets of the CP are highlighted in cyan.

**Extended Data Fig. 7.**
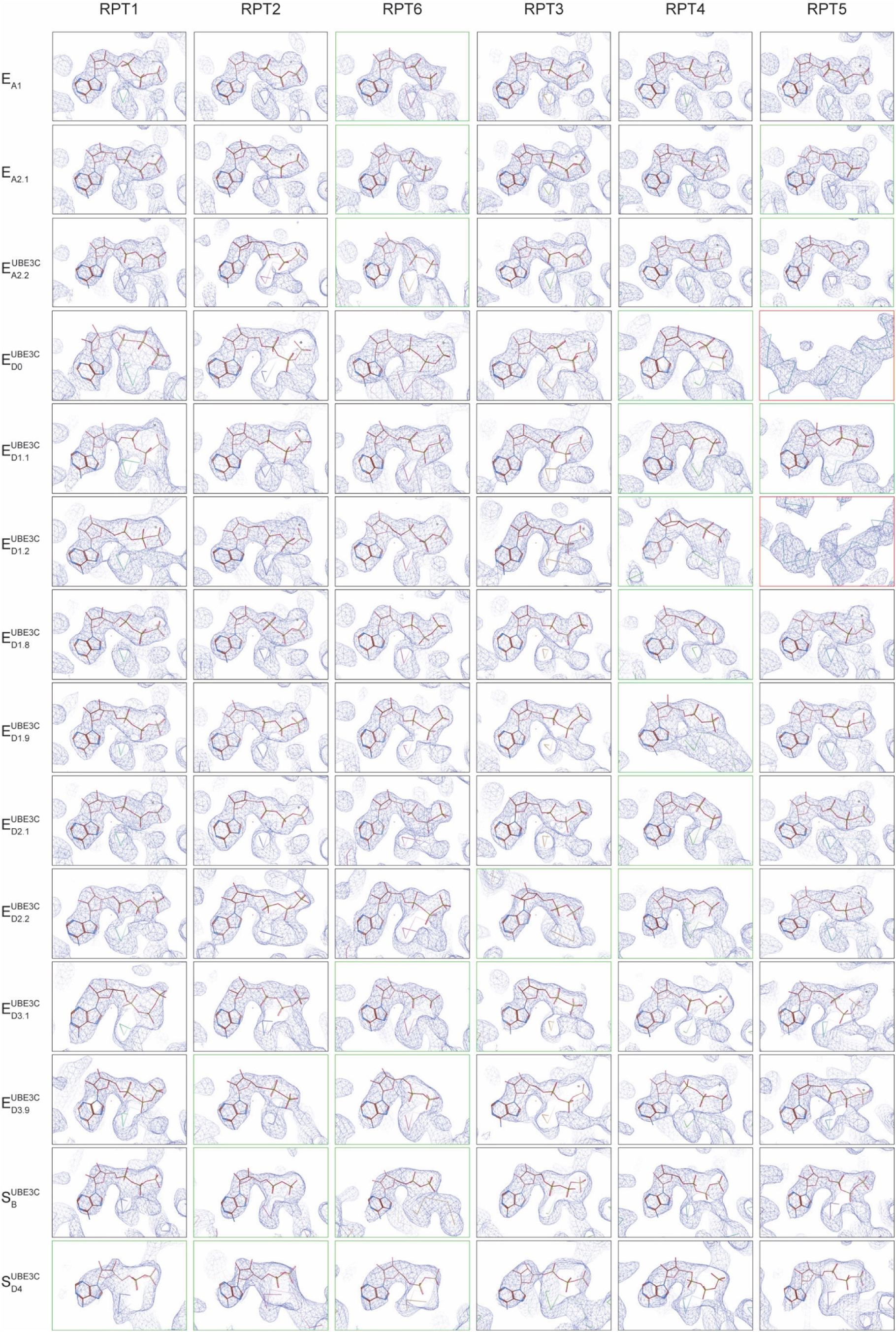
Nucleotide densities in all proteasome states. Comparison of nucleotide densities in 14 proteasome states. The nucleotide densities, fitted with atomic models, are shown as blue mesh representations. All close-up views were directly screen-captured from Coot after atomic modeling into the density maps, without further modification. At the contour level typically used for atomic modeling, the nucleotide densities in the apo-like subunits mostly disappear, though they occasionally appear as partial nucleotide shapes at much lower contour levels. For states with limited local resolutions, nucleotide types are hypothesized based on the densities, the openness of the corresponding nucleotide-binding pockets, and the higher-resolution homologous structural models from states with similar conformations.

**Extended Data Fig. 8.**
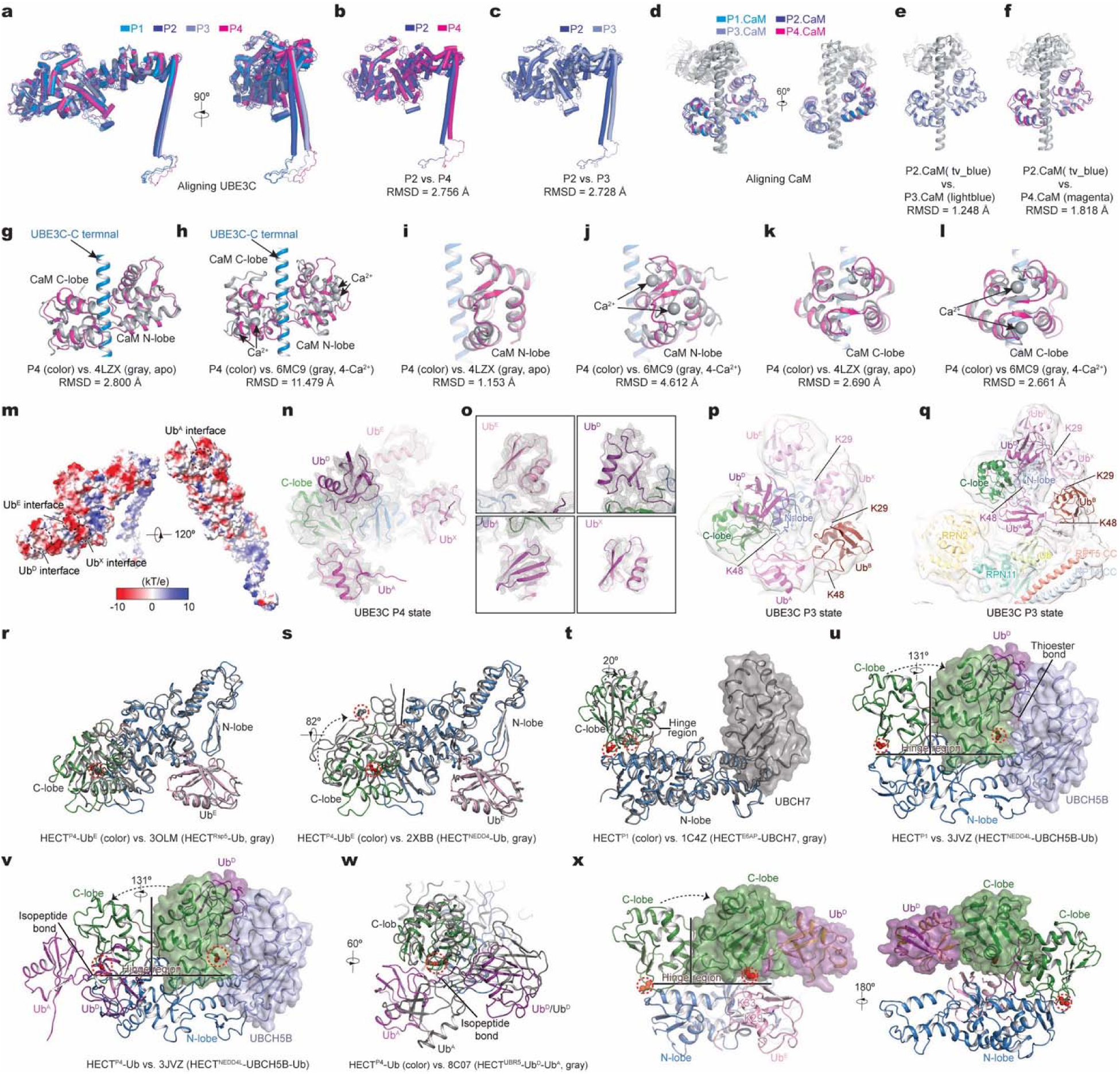
Dynamics of UBE3C and comparison of proteasome-bound UBE3C with other HECT E3 ligases. **a**, Superposition of the UBE3C structures from different states aligned to UBE3C. **b, c**, Structural comparisons of UBE3C between states P2 and P4 (**b**), and between states P2 and P3 (**c**), aligned to UBE3C. **d,** Superposition of calmodulin structures from different UBE3C states aligned to calmodulin. **e, f**, Structural comparisons of calmodulin bound to UBE3C between states P2 and P4 (**e**), and between states P2 and P3 (**f**), aligned to calmodulin. **g-l**, Comparisons of the calmodulin (**g, h**), calmodulin N-lobe (**i, j**), and calmodulin C-lobe (**k, l**) between UBEC3-bound calmodulin and apo calmodulin (PDB ID 4LZX)^82^, and between UBEC3-bound calmodulin and Ca^2+^-bound calmodulin (PDB ID 6MC9)^83^. **m,** Electrostatic surface representation of UBE3C in state P4 showing the acidic or negatively charged surfaces at the four ubiquitin-binding sites designated the A-, D-, E- and X-sites. **n, o,** Cryo-EM densities (grey mesh) of the four well-resolved ubiquitins bound to the UBE3C HECT domain in state P4, superimposed with the atomic model for clear definition of Ub^A^, Ub^D^, Ub^E^ and Ub^X^. Because Ub^X^ exhibits relatively weaker density, a lower contour level and an additional low-pass filter were applied to enhance visibility. **p,** Low-pass filtered cryo-EM map and pseudo-atomic model of ubiquitin chains bound to UBE3C in the P3 state on the E_D_/S_D_-like proteasome, revealing a K29/K48-branched ubiquitin chain topology. **q,** Low-pass filtered cryo-EM map and pseudo-atomic model of ubiquitin chains bound to UBE3C in the P3 state, together with the ubiquitin-bound RPN11 and the RPN2 subunits of the E_A_-like proteasome. **r**, Comparison of HECT-Ub between proteasome bound UBE3C-Ub^E^ and HECT^Rsp5^-Ub (PDB ID 3OLM)^84^, showing a similar Ub-binding conformation. The catalytic Cystine of UBE3C is shown as a sphere and highlighted by a red dashed circle. **s**, Structure comparison of HECT-Ub between proteasome bound UBE3C-Ub^E^ and HECT^NEDD4^-Ub (PDB ID 2XBB)^52^, aligned to the N-lobe of the HECT domain. Both show a similar Ub binding site, but c-lobe of HECT^NEDD4^-Ub is rotated 82°. **t**, Superposition of HECT domains in proteasome bound UBE3C (P1 state) and E6AP-UbcH7 (PDB ID 1C4Z)^85^. When aligned to the N-lobe, the C-lobe of HECT^UBE3C^ is rotated 20° toward UbcH7 around the hinge region. UbcH7 is shown in both cartoon and surface representations. **u, v**, Superpositions of HECT domains in proteasome bound UBE3C (P1 and P4 states) and NEDD4L-UbcH5B-Ub (PDB ID 3JVZ)^49^. Compared with P1 state, the C-lobe of HECT^NEDD4L^ is rotated 131° to approach the transthiolation site conformed by UbcH5B and Ub (**u**). In contrast, the C-lobe in UBE3C P4 state rotates back to face Ub^A^, at which point the Ub^D^ C-terminus is linked to the K48 of Ub^A^ via an isopeptide bond, indicating Ub transfer from E2 to substrate (**v**). NEDD4L-UbcH5B-Ub is shown in both cartoon and surface representations. **w**, Superposition of HECT domains in proteasome bound UBE3C (P4 state) and UBR5-Ub^D^-Ub^A^ (PDB ID 8C07)^50^. **x,** Hypothesized conformation for UBE3C-mediated synthesis of K29/K33 linked Ubiquitin chains. The UBE3C C-lobe and Ub^D^ rotate toward to Ub^E^ such that Ub^D^-G76 approaches K29/K33 of Ub^E^, shown in both cartoon and surface representations.

**Extended Data Fig. 9.**
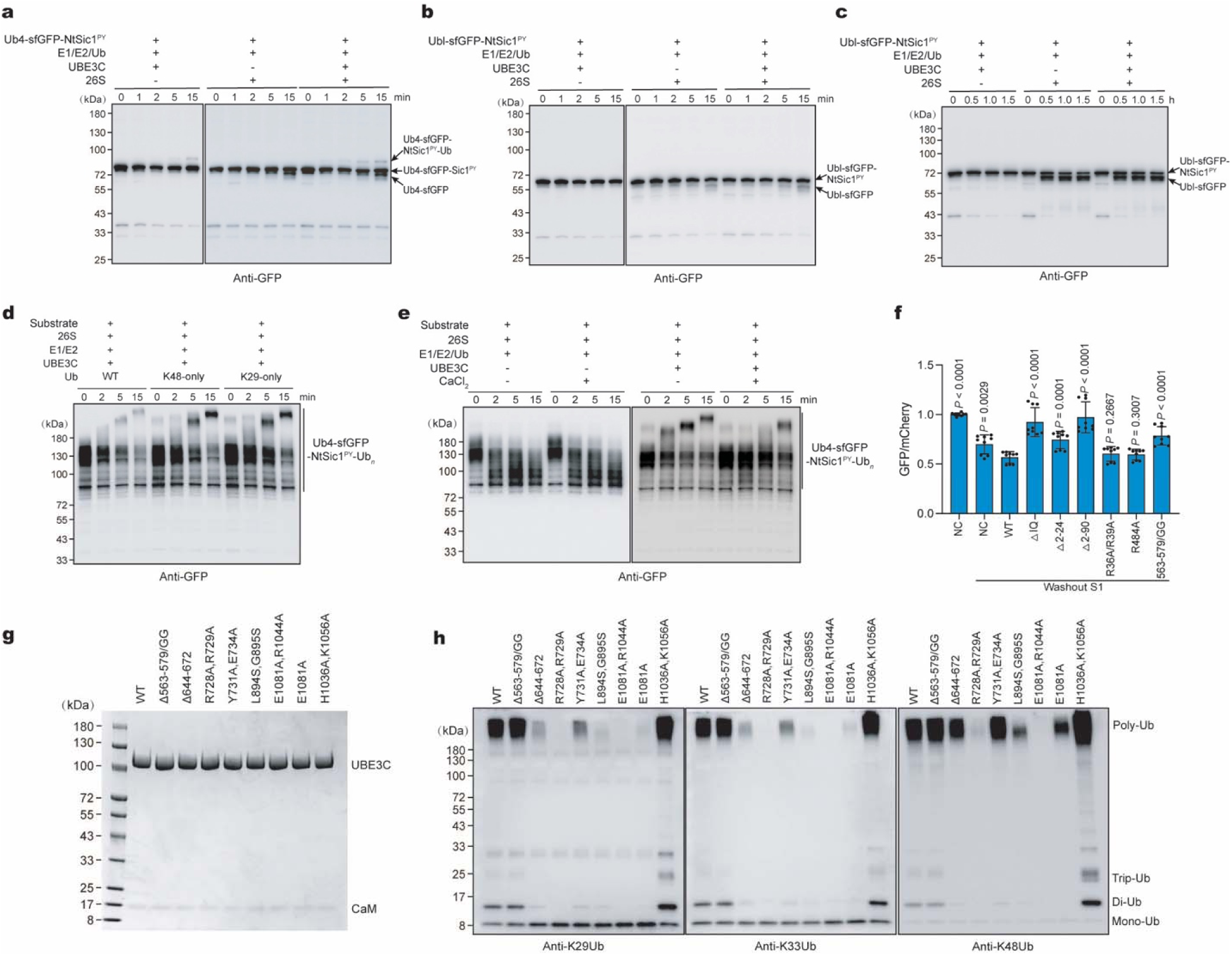
**UBE3C enhanced proteasome processivity and the structure-based site-directed mutagenesis. a-c**, In vitro degradation of a superfolder GFP-based substrate by UBE3C-human proteasome complex in the presence of E1, E2 and Ubiquitin, analyzed by western blot using an anti-GPF antibody. A representative result from three independent experiments is shown. UBE3C has minimal effect on the degradation of Ub4-sfGFP-NtSic1^PY^ **(a)** or Ubl-sfGFP-NtSic1^PY^ **(b),** even following prolonged incubation **(c)**. **d, e,** In vitro degradation of the partially ubiquitylated substrate (Ub4-sfGFP-NtSic1^PY^-K48Ubn) by the UBE3C-human proteasome complex, analyzed by western blot using an anti-GPF antibody. Degradation assays performed in the presence of ubiquitin mutants: K48-only (all lysine residues except K48 mutated to alanine) and K29-only (all lysine residues except K29 mutated to alanine) **(d)**, or in the presence of wildtype ubiquitin and CaCl_2_ **(e)**. Data are representative of three independent experiments. **f**, UBE3C promotes proteasomal processing in cells. UBE3C-knockout HEK293T cells stably expressing DD-sfGFP were transiently transfected with UBE3C variants. sfGFP degradation was assessed by flow cytometry. NC, negative control (pcDNA3.1); WT, the wildtype UBE3C; ΔIQ, IQ motif deletion mutant; Δ2–24 and Δ2–90, N-terminal truncation mutants; 568–579/GG, replacement of residues 568–579 with Gly-Gly. *P*-values were calculated relative to WT using a two-tailed unpaired t-test. Data represent mean ± s.d. from three independent experiments, each experiment includes three replicates. **g**, Purification of UBE3C mutants and analyzed by SDS/PAGE. **h,** In vitro ubiquitylation activity of UBE3C mutants assessed by SDS-PAGE and immunoblotting with antibodies recognizing K29-, K33- and K48-linked ubiquitin chains. Representative results from three independent experiments are shown.

**Extended Data Fig. 10.**
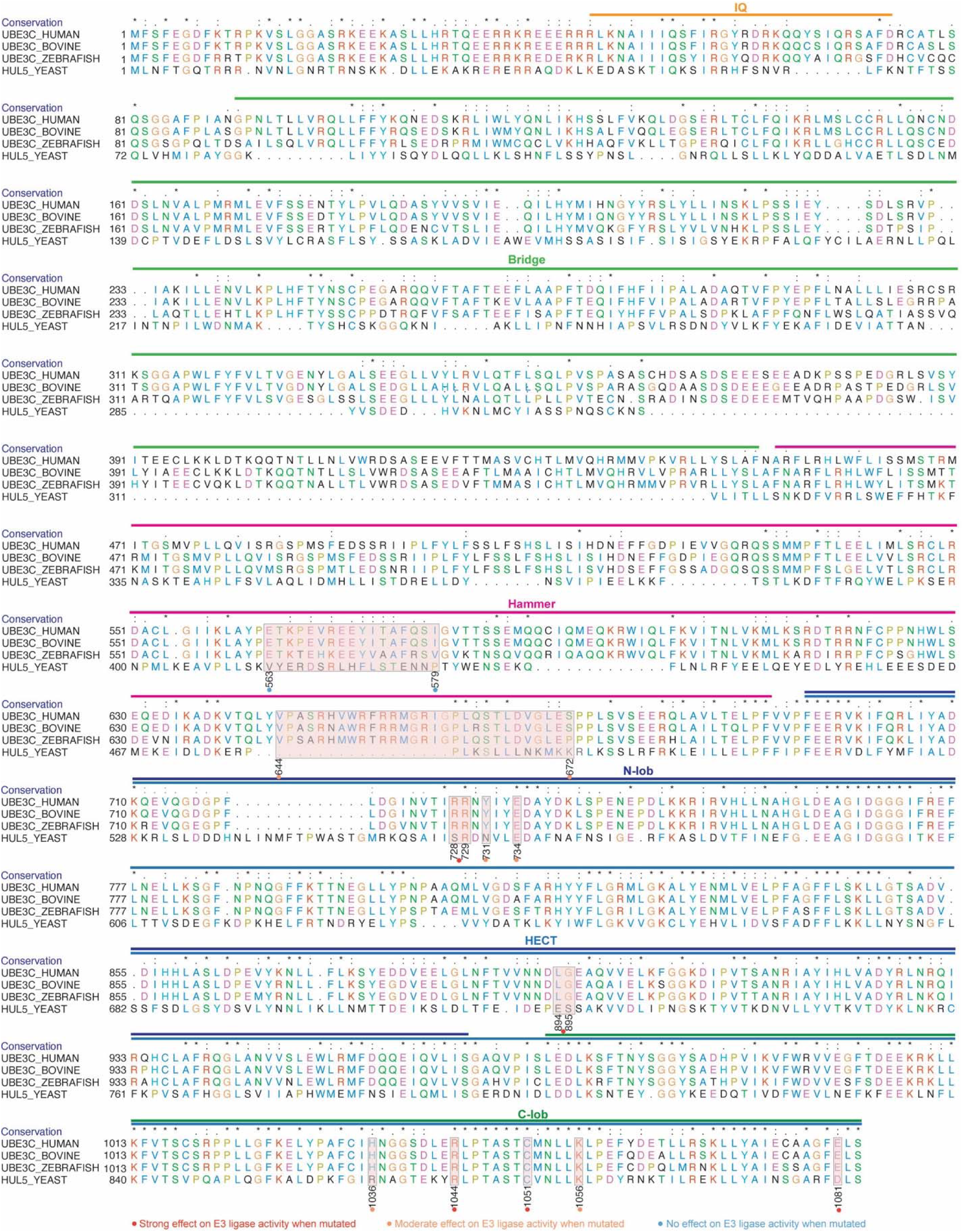
Multiple sequence alignment of UBE3C from four species. Multiple sequence alignment of UBE3C homologs from human, bovine, zebrafish and yeast, performed using UCSF Chimera. Annotations are based on structural and mutational analyses from Fig.4 and Extended Data Fig. 9.

**Extended Data Fig. 11.**
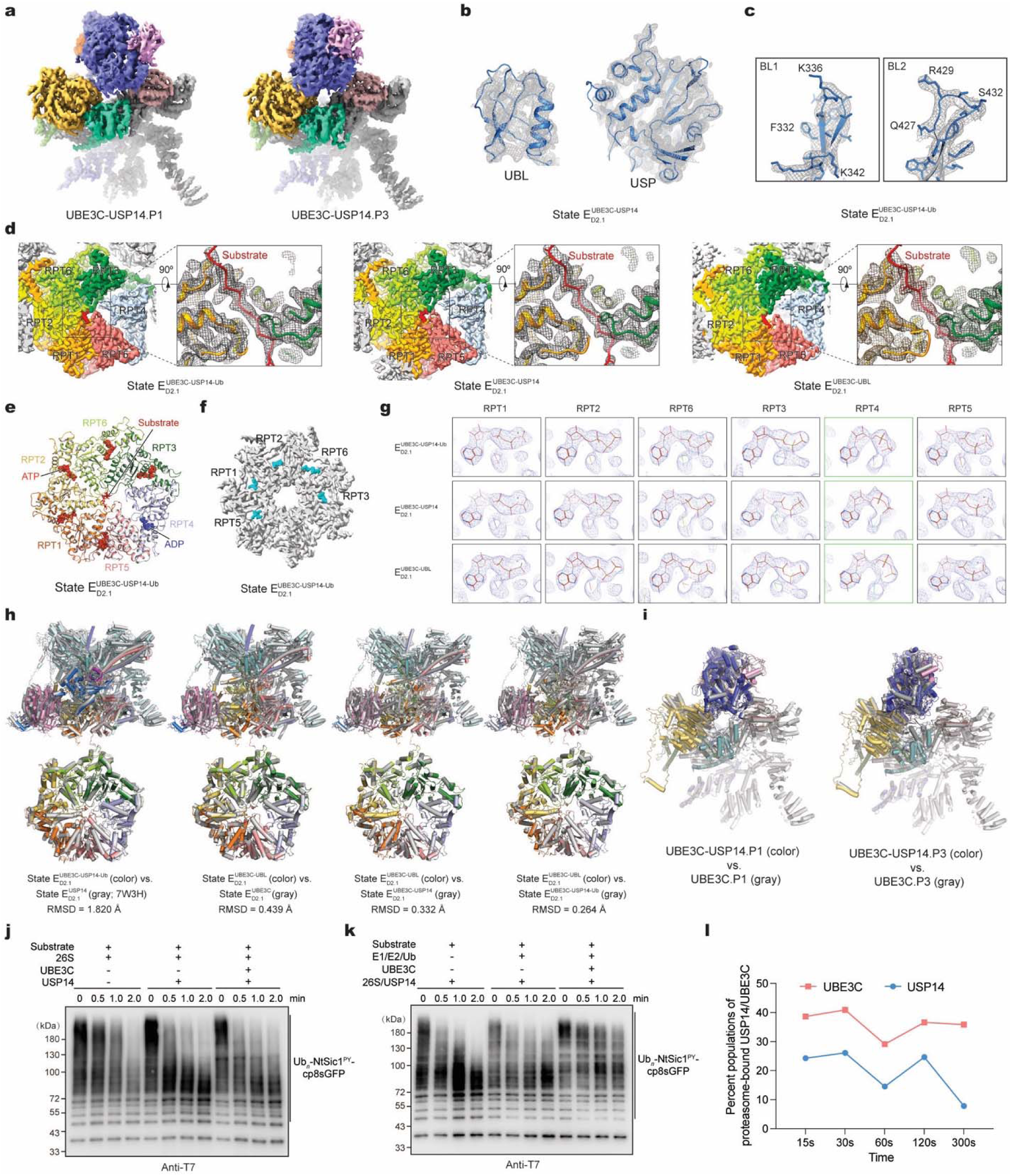
Allosteric antagonism of UBE3C against USP14 at the human proteasome. a, Cryo-EM density maps of two distinct UBE3C conformations (P1 and P3) resolved in UBE3C-USP14-proteasome complexes. b, Cryo-EM densities (mesh) of biquitin-like (UBL) domain and the ubiquitin-specific protease (USP) domain of USP14 in state E_D2.1_^UBE3C-USP14^ **c**, Local cryo-EM densities of the UPS14 blocking loops BL1 and BL2 in E_D2.1_^UBE3C-USP14-Ub^ state, superimposed with the corresponding atomic model (cartoon). **d,** Cryo-EM densities of the AAA-ATPase ring bound to substrate in the UBE3C-USP14-proteasome states E_D2.1_. Top views of the AAA-ATPase ring (left insets) and zoomed-in side views of substrate engagement (right insets) are rendered as surface and mesh, respectively. Substrate densities are modelled as polypeptide backbones and colored in red. **e**, Top views of the AAA-ATPase ring atomic models from UBE3C-USP14-proteasome states E_D2.1_, displayed as cartoon representations. Bound nucleotides are shown as spheres: ADP in blue and ATP in red. **f**, Cryo-EM densities of the RP-CP interface for UBE3C-USP14-proteasome states E_D2.1_, shown in top view. The C-terminal tails of RPT subunits inserted into the α-pockets of the CP are highlighted in cyan. **g,** Comparison of nucleotide densities in UBE3C-USP14-proteasome states E_D2.1_. The nucleotide densities, fitted with atomic models, are shown as blue mesh representations. **h,** Structural comparison of the RP and ATPase structures among UBE3C-USP14-proteasome states, with E_D2.1_^UBE3C^ or with a previously reported cryo-EM structure (PDB ID: 7W3H). The RMSD values for the ATPase are shown below each panel. **i**, Structural comparison of UBE3C P1 and P3 conformations in the UBE3C- proteasome complex versus the UBE3C-USP14-proteasome complex. **j-k,** In vitro degradation of the ubiquitylated substrate by UBE3C-USP14-proteasome complex in the absence of **(j)** or presence of E1, E2 and Ubiquitin **(k)**, analyzed by western blot using an anti-T7 antibody. The substrate, Ub*_n_*-NtSic1^PY^-cp8sfGFP, was generated by partial ubiquitylation of NtSic1^PY^-cp8sfGFP (see Methods). A representative result from three independent experiments is shown. **k,** Time-resolved changes in the particle populations of proteasome-bound UBE3C and USP14 in the proteasome state E_D2.1_, quantified from time-resolved cryo-EM datasets.

**Extended Data Fig. 12.**
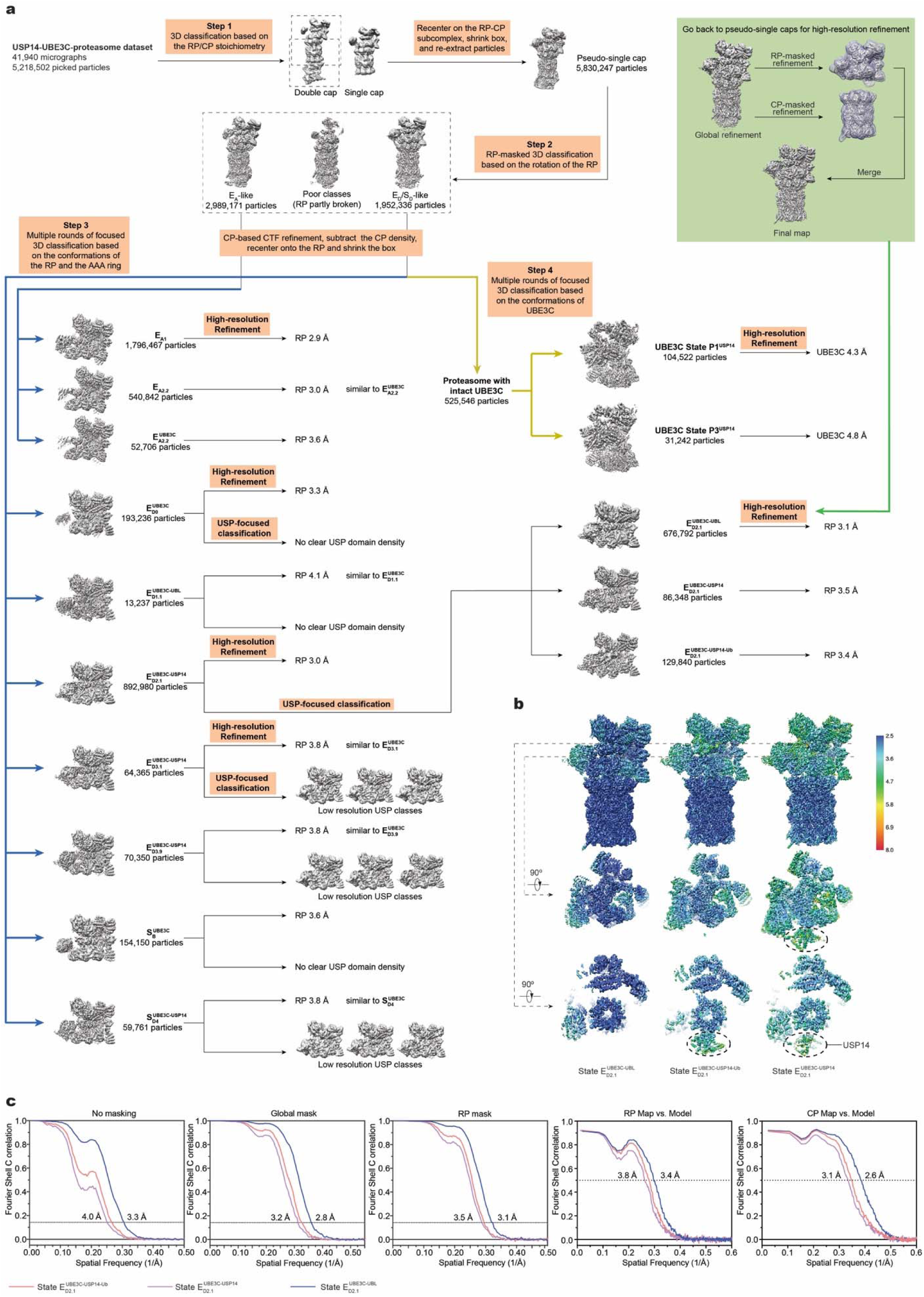
Cryo-EM data processing, reconstructions and resolution measurements for the UBE3C-USP14-proteasome supercomplex. a, Data processing workflow for the USP14-UBE3C-proteasome dataset, which follows a strategy similar to that used for the UBE3C-proteasome dataset (Extended Data Fig. 2a). **b,** Local resolution estimation of the reconstructions of the states E_D2.1_^UBE3C-UBL^, E_D2.1_^UBE3C-USP14^, E_D2.1_^UBE3C-USP14-Ub^. Each state is displayed in two orthogonal views; cross-sections of the top view are shown at the level of the AAA (middle row) and OB (bottom row) domains. **c,** FSC plots for the three states. Plots from left to right show: gold-standard FSC plots of the RP–CP maps calculated without and with masking the separately refined half-maps, gold-standard FSC plot of the RP-masked reconstructions, and model-map FSC plots calculated between each model and its corresponding RP-masked and CP-masked reconstructions.

**Extended Data Table 1.** Cryo-EM data collection, refinement and validation statistics.

| **Proteasome**<br>(UBE3C-proteasome dataset) | $E_{A1}^{UBE3C}$ | $E_{A2.1}^{UBE3C}$ | $E_{A2.2}^{UBE3C}$ | $E_{D0}^{UBE3C}$ | $E_{D1.1}^{UBE3C}$ | $E_{D1.2}^{UBE3C}$ | $E_{D1.8}^{UBE3C}$ |
| --- | --- | --- | --- | --- | --- | --- | --- |
| **Data collection and processing** |  |  |  |  |  |  |  |
| Magnification | 64000 | 64000 | 64000 | 64000 | 64000 | 64000 | 64000 |
| Voltage (kV) | 300 | 300 | 300 | 300 | 300 | 300 | 300 |
| Electron exposure ($e^-/\text{\AA}^2$) | 50 | 50 | 50 | 50 | 50 | 50 | 50 |
| Defocus range ($\mu\text{m}$) | -0.5 to -5.0 | -0.5 to -5.0 | -0.5 to -5.0 | -0.5 to -5.0 | -0.5 to -5.0 | -0.5 to -5.0 | -0.5 to -5.0 |
| Pixel size ($\text{\AA}$) | 0.685 | 0.685 | 0.685 | 0.685 | 0.685 | 0.685 | 0.685 |
| Symmetry imposed | C1 | C1 | C1 | C1 | C1 | C1 | C1 |
| Initial particle images (no.) | 5515349 | 5515349 | 5515349 | 5515349 | 5515349 | 5515349 | 5515349 |
| Final particles images (no.) | 1477218 | 161141 | 160900 | 66299 | 38145 | 47825 | 200634 |
| RP map resolution ($\text{\AA}$) | 2.7 | 3.2 | 3.2 | 3.4 | 3.5 | 3.4 | 3.1 |
| FSC threshold | 0.143 | 0.143 | 0.143 | 0.143 | 0.143 | 0.143 | 0.143 |
| Map resolution range ($\text{\AA}$) | 2.5-8.0 | 2.5-8.0 | 2.5-8.0 | 2.5-8.0 | 2.5-8.0 | 2.5-8.0 | 2.5-8.0 |
| **Refinement** |  |  |  |  |  |  |  |
| Model resolution RP/CP ($\text{\AA}$) | 3.1/2.4 | 3.4/2.8 | 3.4/2.7 | 3.6/2.9 | 3.8/3.1 | 3.5/3.0 | 3.3/2.7 |
| FSC threshold | 0.5 | 0.5 | 0.5 | 0.5 | 0.5 | 0.5 | 0.5 |
| Map sharpening B factor ($\text{\AA}^2$) | -30 | -30 | -30 | 0 | 0 | -30 | -30 |
| **Model composition** |  |  |  |  |  |  |  |
| Non-hydrogen atoms | 105019 | 105078 | 105497 | 106633 | 106480 | 106443 | 106717 |
| Protein residues | 13388 | 13374 | 13424 | 13557 | 13536 | 13533 | 13555 |
| Ligands | 12 | 12 | 11 | 9 | 10 | 10 | 10 |
| **B factors ($\text{\AA}^2$)** |  |  |  |  |  |  |  |
| Protein | 98.76 | 110.65 | 114.46 | 128.61 | 108.85 | 109.4 | 104.64 |
| Ligand | 63.21 | 70.07 | 79.72 | 139.27 | 99.79 | 118.69 | 98.01 |
| **R.m.s. deviations** |  |  |  |  |  |  |  |
| Bond length ($\text{\AA}$) ($\# > 4\sigma$) | 0.004 | 0.004 | 0.005 | 0.004 | 0.003 | 0.005 | 0.005 |
| Bond angles ($^\circ$) ($\# > 4\sigma$) | 0.588 | 0.645 | 0.647 | 0.62 | 0.642 | 0.681 | 0.635 |
| **Validation** |  |  |  |  |  |  |  |
| MolProbity score | 1.52 | 1.58 | 1.59 | 1.58 | 1.62 | 1.65 | 1.59 |
| Clash score | 4.68 | 5.5 | 5.6 | 5.2 | 5.45 | 5.64 | 5.19 |
| Rotamer outliers (%) | 0.15 | 0.1 | 0.11 | 0.06 | 0.11 | 0.06 | 0.15 |
| **Ramachandran plot** |  |  |  |  |  |  |  |
| Outliers (%) | 0.09 | 0.05 | 0.05 | 0.06 | 0.09 | 0.15 | 0.15 |
| Allowed (%) | 3.98 | 4.11 | 4.11 | 4.24 | 4.54 | 4.80 | 4.37 |
| Favored (%) | 95.93 | 95.84 | 95.84 | 95.70 | 95.37 | 95.05 | 95.49 |

| **Proteasome**<br>(UBE3C-proteasome dataset) | $E_{D1.9}^{UBE3C}$ | $E_{D2.1}^{UBE3C}$ | $E_{D2.2}^{UBE3C}$ | $E_{D3.1}^{UBE3C}$ | $E_{D3.9}^{UBE3C}$ | $S_B^{UBE3C}$ | $S_{D4}^{UBE3C}$ |
| --- | --- | --- | --- | --- | --- | --- | --- |
| **Data collection and processing** |  |  |  |  |  |  |  |
| Magnification | 64000 | 64000 | 64000 | 64000 | 64000 | 64000 | 64000 |
| Voltage (kV) | 300 | 300 | 300 | 300 | 300 | 300 | 300 |
| Electron exposure ($e^-/\text{\AA}^2$) | 50 | 50 | 50 | 50 | 50 | 50 | 50 |
| Defocus range ($\mu\text{m}$) | -0.5 to -5.0 | -0.5 to -5.0 | -0.5 to -5.0 | -0.5 to -5.0 | -0.5 to -5.0 | -0.5 to -5.0 | -0.5 to -5.0 |
| Pixel size ($\text{\AA}$) | 0.685 | 0.685 | 0.685 | 0.685 | 0.685 | 0.685 | 0.685 |
| Symmetry imposed | C1 | C1 | C1 | C1 | C1 | C1 | C1 |
| Initial particle images (no.) | 5515349 | 5515349 | 5515349 | 5515349 | 5515349 | 5515349 | 5515349 |
| Final particles images (no.) | 196654 | 669095 | 58022 | 54371 | 129985 | 24115 | 15183 |
| RP map resolution ($\text{\AA}$) | 3.2 | 2.9 | 3.5 | 3.5 | 3.3 | 3.8 | 4.1 |
| FSC threshold | 0.143 | 0.143 | 0.143 | 0.143 | 0.143 | 0.143 | 0.143 |
| Map resolution range ($\text{\AA}$) | 2.5-8.0 | 2.5-8.0 | 2.5-8.0 | 2.5-8.0 | 2.5-8.0 | 2.5-8.0 | 2.5-8.0 |
| **Refinement** |  |  |  |  |  |  |  |
| Model resolution RP/CP ($\text{\AA}$) | 3.4/2.7 | 3.1/2.5 | 3.9/3.0 | 3.8/3.0 | 3.5/2.8 | 4.2/3.2 | 4.4/3.3 |
| FSC threshold | 0.5 | 0.5 | 0.5 | 0.5 | 0.5 | 0.5 | 0.5 |
| Map sharpening B factor ($\text{\AA}^2$) | -30 | -30 | -30 | -30 | -30 | -30 | 0 |
| **Model composition** |  |  |  |  |  |  |  |
| Non-hydrogen atoms | 106717 | 106640 | 106636 | 106616 | 106622 | 106302 | 106435 |
| Protein residues | 13555 | 13544 | 13556 | 13550 | 13550 | 13520 | 13543 |
| Ligands | 10 | 10 | 9 | 9 | 10 | 7 | 7 |
| B factors ($\text{\AA}^2$) | | | | | | | |
| Protein | 112.51 | 108.24 | 100.89 | 99.21 | 119.73 | 96.96 | 149.37 |
| Ligand | 104.46 | 95.96 | 87.67 | 90.43 | 134.67 | 83.05 | 133.28 |
| R.m.s. deviations |  |  |  |  |  |  |  |
| Bond length ($\text{\AA}$) ($\# > 4 \sigma$) | 0.004 | 0.005 | 0.003 | 0.004 | 0.003 | 0.005 | 0.004 |
| Bond angles ($^\circ$) ($\# > 4 \sigma$) | 0.675 | 0.62 | 0.586 | 0.642 | 0.669 | 0.67 | 0.65 |
| Validation |  |  |  |  |  |  |  |
| MolProbity score | 1.62 | 1.58 | 1.6 | 1.69 | 1.62 | 1.66 | 1.65 |
| Clash score | 5.51 | 5.08 | 5.1 | 5.88 | 5.58 | 5.67 | 5.86 |
| Rotamer outliers (%) | 0.12 | 0.08 | 0.02 | 0.16 | 0.08 | 0.04 | 0.07 |
| Ramachandran plot |  |  |  |  |  |  |  |
| Outliers (%) | 0.11 | 0.09 | 0.07 | 0.18 | 0.11 | 0.10 | 0.07 |
| Allowed (%) | 4.59 | 4.43 | 4.69 | 5.13 | 4.40 | 5.00 | 4.69 |
| Favored (%) | 95.30 | 95.48 | 95.24 | 94.69 | 95.49 | 94.90 | 95.23 |

| **Proteasome**<br>(USP14-UBE3C-proteasome dataset) | $E_{D2.1}^{\text{UBE3C-UBL}}$ | $E_{D2.1}^{\text{UBE3C-USP14}}$ | $E_{D2.1}^{\text{UBE3C-USP14-Ub}}$ |
| --- | --- | --- | --- |
| **Data collection and processing** |  |  |  |
| Magnification | 64000 | 64000 | 64000 |
| Voltage (kV) | 300 | 300 | 300 |
| Electron exposure ($e^-/\text{\AA}^2$) | 50 | 50 | 50 |
| Defocus range ($\mu\text{m}$) | -0.5 to -5.0 | -0.5 to -5.0 | -0.5 to -5.0 |
| Pixel size ($\text{\AA}$) | 0.685 | 0.685 | 0.685 |
| Symmetry imposed | C1 | C1 | C1 |
| Initial particle images (no.) | 5218502 | 5218502 | 5218502 |
| Final particles images (no.) | 676792 | 86348 | 129840 |
| RP map resolution ($\text{\AA}$) | 3.1 | 3.5 | 3.4 |
| FSC threshold | 0.143 | 0.143 | 0.143 |
| Map resolution range ($\text{\AA}$) | 2.5-8.0 | 2.5-8.0 | 2.5-8.0 |
| **Refinement** |  |  |  |
| Model resolution RP/CP ($\text{\AA}$) | 3.4/2.6 | 3.8/3.1 | 3.6/2.9 |
| FSC threshold | 0.5 | 0.5 | 0.5 |
| Map sharpening B factor ($\text{\AA}^2$) | -30 | -30 | -30 |
| Model composition |  |  |  |
| Non-hydrogen atoms | 107043 | 110230 | 110811 |
| Protein residues | 13601 | 13995 | 14071 |
| Ligands | 9 | 9 | 9 |
| B factors ($\text{\AA}^2$) | | | |
| Protein | 115.75 | 102.41 | 111.15 |
| Ligand | 104.86 | 98.43 | 99.39 |
| R.m.s. deviations |  |  |  |
| Bond length ($\text{\AA}$) ($\# > 4 \sigma$) | 0.005 | 0.003 | 0.003 |
| Bond angles ($^\circ$) ($\# > 4 \sigma$) | 0.631 | 0.590 | 0.609 |
| Validation |  |  |  |
| MolProbity score | 1.53 | 1.52 | 1.55 |
| Clash score | 4.49 | 4.42 | 4.82 |
| Rotamer outliers (%) | 0.09 | 0.06 | 0.05 |
| Ramachandran plot |  |  |  |
| Outliers (%) | 0.09 | 0.09 | 0.09 |
| Allowed (%) | 4.36 | 4.20 | 4.28 |
| Favored (%) | 95.55 | 95.72 | 95.62 |

| **UBE3C**<br>(UBE3C-proteasome dataset) | P1 | P2 | P3 | P4 |
| --- | --- | --- | --- | --- |
| **Data collection and processing** |  |  |  |  |
| Magnification | 64000 | 64000 | 64000 | 64000 |
| Voltage (kV) | 300 | 300 | 300 | 300 |
| Electron exposure (e<sup>-</sup>/Å<sup>2</sup>) | 50 | 50 | 50 | 50 |
| Defocus range (μm) | -0.5 to -5.0 | -0.5 to -5.0 | -0.5 to -5.0 | -0.5 to -5.0 |
| Pixel size (Å) | 0.685 | 0.685 | 0.685 | 0.685 |
| Symmetry imposed | C1 | C1 | C1 | C1 |
| Initial particle images (no.) | 5515349 | 5515349 | 5515349 | 5515349 |
| Final particles images (no.) | 284038 | 99398 | 63572 | 91293 |
| UBE3C map resolution (Å) | 3.8 | 4.2 | 4.4 | 3.8 |
| FSC threshold | 0.143 | 0.143 | 0.143 | 0.143 |
| Map resolution range (Å) | 3.0-8.0 | 3.0-8.0 | 3.0-8.0 | 3.0-8.0 |
| **Refinement** |  |  |  |  |
| Model resolution Lid-UBE3C (Å) | 3.4 | 3.5 | 3.8 | 3.5 |
| FSC threshold | 0.5 | 0.5 | 0.5 | 0.5 |
| Map sharpening B factor (Å<sup>2</sup>) | -70 | -50 | -70 | -75 |
| Model composition |  |  |  |  |
| Non-hydrogen atoms | 41007 | 41130 | 42227 | 42177 |
| Protein residues | 5113 | 5130 | 5268 | 5261 |
| Ligands | 1 | 1 | 1 | 1 |
| B factors (Å<sup>2</sup>) |  |  |  |  |
| Protein | 146.73 | 146.72 | 160.93 | 64.92 |
| Ligand | 140.69 | 134.14 | 153.8 | 129.9 |
| R.m.s. deviations |  |  |  |  |
| Bond length (Å) (# > 4 σ ) | 0.003 | 0.004 | 0.004 | 0.003 |
| Bond angles (°) (# > 4 σ ) | 0.631 | 0.68 | 0.659 | 0.648 |
| Validation |  |  |  |  |
| MolProbity score | 1.72 | 1.84 | 1.8 | 1.73 |
| Clash score | 5.61 | 6.6 | 6.17 | 5.48 |
| Rotamer outliers (%) | 0.2 | 0.22 | 0.26 | 0.15 |
| Ramachandran plot |  |  |  |  |
| Outliers (%) | 0.22 | 0.26 | 0.23 | 0.23 |
| Allowed (%) | 6.07 | 7.34 | 7.04 | 6.55 |
| Favored (%) | 93.71 | 92.41 | 92.73 | 93.22 |

**Extended Data Table 2:** Summary of key structural features of the UBE3C-bound proteasome in different states.

|  | UBE3C binding | CP gate state | Locations of substrate densities | Pore-loop staircase contacting substrate (from top to bottom) | RPT subunits with substrate-disengaged pore-1 loop | RPT subunits in ADP-bound state | RPT subunits in apo-like state | USP14 binding site |
| --- | --- | --- | --- | --- | --- | --- | --- | --- |
| E<sup>UBE3C</sup><sub>A1</sub> | No | Closed | None | None | - | RPT6 | None | - |
| E<sup>UBE3C</sup><sub>A2.1</sub> | No | Closed | None | None | - | RPT6 | None | - |
| E<sup>UBE3C</sup><sub>A2.2</sub> | Yes | Closed | None | None | - | RPT6 | None | - |
| E<sup>UBE3C</sup><sub>D0</sub> | Yes | Open | AAA channel | RPT2, RPT6, RPT3, RPT4, RPT5 | RPT1 | RPT4 | RPT5 | - |
| E<sup>UBE3C</sup><sub>D1.1</sub> | Yes | Open | AAA channel | RPT2, RPT6, RPT3, RPT4 | RPT1, RPT5 | RPT4, RPT5 | None | - |
| E<sup>UBE3C</sup><sub>D1.2</sub> | Yes | Open | AAA channel | RPT1, RPT2, RPT6, RPT3, RPT4 | RPT5 | RPT4 | RPT5 | - |
| E<sup>UBE3C</sup><sub>D1.8</sub> | Yes | Open | AAA channel | RPT1, RPT2, RPT6, RPT3, RPT4 | RPT5 | RPT4 | None | - |
| E<sup>UBE3C</sup><sub>D1.9</sub> | Yes | Open | AAA channel | RPT5, RPT1, RPT2, RPT6, RPT3, | None | RPT4 | None | - |
|  |  |  |  | RPT4 |  |  |  |  |
| $E_{D2.1}^{UBE3C}$ | Yes | Open | AAA channel | RPT5,<br>RPT1,<br>RPT2,<br>RPT6,<br>RPT3 | RPT4 | RPT4 | None | - |
| $E_{D2.2}^{UBE3C}$ | Yes | Open | AAA channel | RPT5,<br>RPT1,<br>RPT2,<br>RPT6,<br>RPT3 | RPT4 | RPT3, RPT4 | None | - |
| $E_{D3.1}^{UBE3C}$ | Yes | Open | AAA channel | RPT4,<br>RPT5,<br>RPT1,<br>RPT2,<br>RPT6 | RPT3 | RPT6, RPT3 | None | - |
| $E_{D3.9}^{UBE3C}$ | Yes | Open | AAA channel | RPT3,<br>RPT4,<br>RPT5,<br>RPT1,<br>RPT2,<br>RPT6 | None | RPT2, RPT6 | None | - |
| $S_B^{UBE3C}$ | Yes | Closed | None | None | - | RPT2, RPT6 | None | - |
| $S_{D4}^{UBE3C}$ | Yes | Open | None | None | - | RPT1, RPT2 | None | - |
| $E_{D2.1}^{UBE3C-UBL}$ | Yes | Open | AAA channel | RPT5,<br>RPT1,<br>RPT2,<br>RPT6,<br>RPT3 | RPT4 | RPT4 | None | RPN1<br>T2 site<br>(UBL) |
| $E_{D2.1}^{UBE3C-USP14}$ | Yes | Open | AAA channel | RPT5,<br>RPT1,<br>RPT2,<br>RPT6,<br>RPT3 | RPT4 | RPT4 | None | RPN1<br>T2,<br>RPT1<br>AAA |
| $E_{D2.1}^{UBE3C-USP14-Ub}$ | Yes | Open | AAA channel | RPT5,<br>RPT1,<br>RPT2, | RPT4 | RPT4 | None | RPN1<br>T2,<br>RPT1 |
|  |  |  |  | RPT6,<br>RPT3 |  |  |  | OB |

